# New tools for carbohydrate sulphation analysis: Heparan Sulphate 2-*O*-sulphotranserase (HS2ST) is a target for small molecule protein kinase inhibitors

**DOI:** 10.1101/296533

**Authors:** Dominic P Byrne, Yong Li, Krithika Ramakrishnan, Igor L Barsukov, Edwin A Yates, Claire E Eyers, Dulcé Papy-Garcia, Sandrine Chantepie, Vijayakanth Pagadala, Jian Liu, Carrow Wells, David H Drewry, William J Zuercher, Neil G Berry, David G Fernig, Patrick A Eyers

**Affiliations:** Department of Biochemistry, Institute of Integrative Biology, University of Liverpool, L69 7ZB, UK.; Centre for Proteome Research, Institute of Integrative Biology, University of Liverpool, L69 7ZB, UK.; Laboratory CRRET CNRS 9215, Université Paris-Est, CRRET (EA 4397/ERL CNRS 9215), UPEC, F-94010, Créteil, France.; Glycan Therapeutics, 617 Hutton Street, Raleigh, NC 27606, USA.; UNC Eshelman School of Pharmacy, University of North Carolina at Chapel Hill, Chapel Hill, NC, 27599, USA.; Structural Genomics Consortium, UNC Eshelman School of Pharmacy, University of North Carolina at Chapel Hill, Chapel Hill, NC, 27599, USA.; Lineberger Comprehensive Cancer Center, University of North Carolina at Chapel Hill, Chapel Hill, NC 27599, USA.; Department of Chemistry, University of Liverpool, L69 7ZD, UK

**Keywords:** HS2ST, PAPS, glycan, substrate PAPS, screening, enzyme, kinase, inhibitor

## Abstract

Sulphation of carbohydrate residues occurs on a variety of glycans destined for secretion, and this modification is essential for efficient matrix-based signal transduction. Heparan sulphate (HS) glycosaminoglycans control physiological functions ranging from blood coagulation to cell proliferation. HS biosynthesis involves membrane-bound Golgi sulphotransferases, including heparan sulphate 2-*O*-sulphotransferase (HS2ST), which transfers sulphate from the co-factor PAPS (3’-phosphoadenosine 5’-phosphosulphate) to the 2-*O* position of α-L-iduronate in the maturing oligosaccharide chain. The current lack of simple non-radioactive enzyme assays that can be used to quantify the levels of carbohydrate sulphation hampers kinetic analysis of this process and the discovery of HS2ST inhibitors. In this paper, we describe a new procedure for thermal shift analysis of purified HS2ST. Using this approach, we quantify HS2ST-catalyzed oligosaccharide sulphation using a novel synthetic fluorescent substrate and screen the Published Kinase Inhibitor Set (PKIS), to evaluate compounds that inhibit catalysis. We report the susceptibility of HS2ST to a variety of cell permeable compounds *in vitro*, including polyanionic polar molecules, the protein kinase inhibitor rottlerin and oxindole-based RAF kinase inhibitors. In a related study, published back-to-back with this article, we demonstrate that Tyrosyl Protein Sulpho Tranferases (TPSTs) are also inhibited by a variety of protein kinase inhibitors. We propose that appropriately validated small molecule compounds could become new tools for rapid inhibition of glycan (and protein) sulphation in cells, and that protein kinase inhibitors might be repurposed or redesigned for the specific inhibition of HS2ST.

**SUMMARY STATEMENT:** We report that HS2ST, which is a PAPS-dependent glycan sulphotransferase, can be assayed using a variety of novel biochemical procedures, including a non-radioactive enzyme-based assay that detects glycan substrate sulphation in real time. HS2ST activity can be inhibited by different classes of compounds, including known protein kinase inhibitors, suggesting new approaches to evaluate the roles of HS2ST-dependent sulphation with small molecules in cells.

## INTRODUCTION

Biological sulphation is a widespread reversible covalent modification found throughout nature [1]. The regulated sulphation of saccharides is critical for cellular signalling, including regulatory interactions between extracellular glycoproteins that control signal transduction and high-affinity interactions between different cellular surfaces [2]. In addition to providing mechanical strength, the sulphate-rich extracellular matrix also represents a hub for sulphation-based communication through growth factor signalling [3]. For example, FGF-receptor interactions and intracellular signaling to the ERK pathway are blunted in the absence of appropriate 2-*O* sulphation driven by Heparan Sulphate (HS)-modifying enzymes [4–9], while sulphation of the tetrasaccharide Sialyl Lewis^X^ antigen on glycolipids controls leukocyte adhesion to the endothelium during inflammation [10, 11]. Inappropriate glycan sulphation can therefore underlie aspects of abnormal signalling, infection, inflammation and, increasingly, human neuropathies [12], suggesting that targeting of carbohydrate sulphation dynamics using small molecule enzyme inhibitors remains a priority in both basic and translational research [13]. Indeed, the current limited chemical toolbox to rapidly modify and study glycan sulphation is based around small molecule inhibitors of Sulphatase-2 (Sulf-2), such as OKN-007 [14] or heparanase inhibitors and HS mimics, including roneparstat and PG545, which have been employed for basic and clinical investigation [15].

Glycan sulphotransferases (STs) can be classified into several families depending upon the positional substrate specificity of enzymes for their respective sugar substrates [16, 17]. Heparan Sulphate 2*-O*-sulphotransferase (HS2ST) is required for the generation of Heparan Sulphate (HS), which is an abundant unbranched extracellular glycosaminoglycan with key roles in a range of physiological functions, most notably growth-factor dependent signalling related to development, cell migration and inflammation [18]. HS2ST is a transmembrane protein whose catalytic domain faces into the lumen of the Golgi compartment, and catalyses the sulphation of iduronic acid and, to a lesser extent β-D-glucouronate (GlcA), during the enzymatic assembly of secretory proteoglycans such as HS [18, 19]. HS2ST transfers the sulpho-moiety from PAPS (3’-phosphoadenosine 5’-phosphosulphate) sulphate donor to the C2 hydroxyl of IdoA that lies adjacent to an *N*-sulphated glucosamine residue, generating a 2-*O*-sulphated saccharide unit [20–22]. Removal of the sulphate by endosulphatases such as Sulf-2, or more general HS processing by heparanase, also contributes to the complex physiological patterns of carbohydrate editing found *in vivo* [23].

The analysis of murine models lacking HS2ST reveals central roles for 2-*O*-sulphated HS in kidney development and neuronal function, and for signalling through Wnt and FGF-dependent pathways [8, 18, 24–26]. However, in order to carefully control and examine the dynamics and structural heterogeneity of 2-*O* sulphation patterns in HS, which are the consequences of nontemplate-based synthesis of HS and complex dynamic sulphation patterns, new small molecule approaches for the direct, reversible, inhibition of sulphotransferase enzymes are urgently required. In particular, these need to be deployed using chemical biology strategies to overcome deficiencies associated with genetic disruption approaches relevant to development and/or compensatory glycosylation or signalling mechanisms [27].

Mechanistic parallels between the enzymatic pathway of biological sulphation by sulphotransferases [28] and phosphorylation by protein kinases [29] are apparent, since both enzyme classes transfer charged chemical units from an adenine-based nucleotide co-factor to a (usually) polymeric acceptor structure. The biological analysis of protein kinases, which are thought to employ a similar ‘in-line’ enzyme reaction as the 2-*O* sulphotransferases [28] when transferring phosphate to peptide targets [30], has been revolutionised by the synthesis and wide availability of small molecule inhibitors [31]. Many of these compounds were originally discovered in screens with ATP-competitive inhibitor libraries using oncology-associated target enzymes [32]. Protein kinases have proven to be exceptional targets for the development of therapeutic agents in humans, and dozens of kinase inhibitors have been approved, or will soon be approved, for cancer and anti-inflammatory indications [33]. To help diversify and accelerate this process, validated open-source panels of such inhibitors, such as the Public Kinase Inhibitor Set (PKIS), have been assembled for screening purposes, constituting a variety of chemotypes for unbiased small molecule inhibitor discovery, which can be applied to a diverse range of protein targets [34].

The analysis of carbohydrate sulphation currently relies heavily on genetic, biophysical (NMR) and combinatorial organic chemistry and enzymatic analysis, with only a handful of low-affinity inhibitors of carbohydrate sulphotransferases ever having been disclosed [13, 35]. More recently, a relatively potent inhibitor of the related Type IV aryl sulphotransferase [36] and much lower affinity oestrogen sulphotransferases inhibitors [37–39] were reported. Due to a lack of any selective chemical tool compounds, cellular glycan sulphation remains highly understudied, relying on non-specific cellular methods such as chlorate exposure [40], and the field remains ripe for technological innovation and new chemical biology approaches. Early attempts to discover such molecules amongst small, relatively unfocussed, kinase-based libraries led to the discovery of low-affinity purine and tyrphostin-based inhibitory compounds, which are well-established chemical classes of protein kinase inhibitor [35]. This raises the question as to whether PAPS-dependent sulphotransferases are general inhibitory targets for new or repurposed small molecules that target nucleotide-binding sites, especially broader families of compounds originally developed as protein kinase inhibitors. However, the low throughput nature of radioactive (^35^S-PAPS) TLC or HPLC-based assays typically used for sulphotransferase analysis [35, 41], and the low potency of current hits, argues for new approaches to assay and screen a diverse selection of chemical libraries.

In this paper, we describe new *in vitro* methods for assaying recombinant HS2ST, one of which employs a fluorescent-based detection system with a hexasaccharide substrate. PAPS-dependent sulphation of the substrate at the 2-*O* position of the IdoA residue leads to a change in substrate chemical properties, which can be detected as a real-time mobility shift in a high-throughput microfluidic assay format originally developed for the analysis of peptide phosphorylation [42]. We exploit this assay alongside differential scanning fluorimetry (DSF) to screen a small molecule PKIS library, characterising HS2ST susceptibility towards a variety of cell permeable ligands, including polyanionic chemicals, the promiscuous protein kinase inhibitor rottlerin and a family of oxindole-based inhibitors of the proto-oncogene RAF. We propose that appropriately validated small molecule ligands might become invaluable probes for rapid cellular inhibition of HS2STs, and that further iteration could lead to the synthesis (or repurposing) of small molecules, including compound classes currently employed as kinase inhibitors, to probe cellular HS2ST function.

## EXPERIMENTAL: MATERIALS AND METHODS

### Chemicals and Compounds

Heparin or oligomeric saccharide standards, termed dp2-dp12 [43], or polymeric sulphated heparin-derivatives (Table 1) were synthesised in-house as previously described [44]. *N*-sulphated, fluorescein-tagged hexasaccharide glycan substrate (GlcNS-GlcA-GlcNS-IdoA-GlcNS-GlcA fluorescein, where S=sulphation) containing either L-IdoA or GlcA residues at the third residue from the reducing end (to which a linker and the fluorophore were conjugated) were both purchased from GLYCAN therapeutics. All standard laboratory biochemicals, were purchased from either Melford or Sigma, and were of the highest analytical quality. PAPS (adenosine 3’-phosphate 5’-phosphosulphate, lithium salt hydrate, APS (adenosine 5’-phosphosulphate, sodium salt), PAP (adenosine 3’-5’-diphosphate, disodium salt), CoA (coenzymeA, sodium salt) dephosphoCoA (3’-dephosphoCoA, sodium salt hydrate), ATP (adenosine 5’-triphosphate, disodium salt hydrate) ADP (adenosine 5’-diphosphate, disodium salt), AMP (adenosine 5’-monophosphate, sodium salt), GTP (guanosine 5’-triphosphate, sodium salt hydrate), GDP (guanosine 5’-diphosphate, sodium salt) or cAMP (adenosine 3’,5’-cyclic monophosphate, sodium salt) were all purchased from Sigma and stored at -80°C to minimise degradation. Rottlerin, suramin, aurintricarboxylic acid and all named kinase inhibitors were purchased from Sigma, BD laboratories, Selleck or Tocris.

**Table 1.**
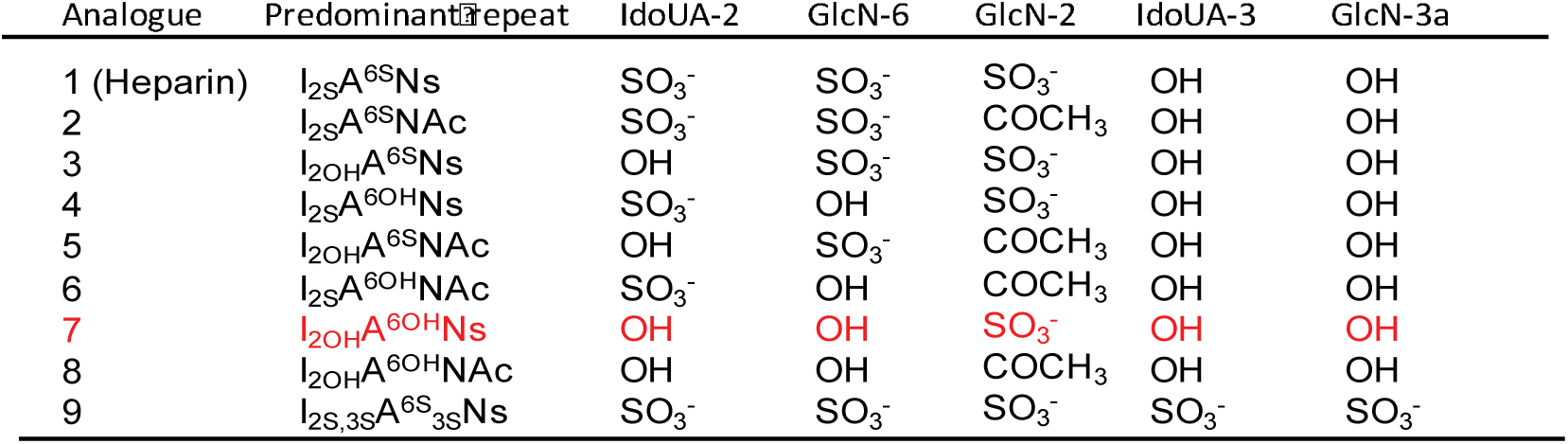
Predominant substitution patterns of differentially-sulphated heparin derivatives described in this study.

### Cloning, recombinant protein production and SDS-PAGE

Chicken HS2ST (isoform 1), which exhibits ~92% identity to human HS2ST, was a kind gift from Dr Lars Pedersen (NIH, USA), and was expressed in the Rosetta-gami (DE3) strain of *E. coli* from a modified pMAL-c2x plasmid encoding an N-terminal Maltose Binding Protein (MBP) affinity tag. Trimeric recombinant HS2ST1 enzyme was partially purified using immobilised amylose affinity chromatography directly from the cleared bacterial extract, essentially as described previously [28]. MBP-HS2ST was eluted with maltose and further purified by SEC using a HiLoad 16/600 Superdex 200 column (GE Healthcare), which was equilibrated in 50 mM Tris–HCl, pH 7.4, 100 mM NaCl, 10% (v/v) glycerol and 1 mM DTT. Prior to analysis, purified proteins were snap frozen in liquid nitrogen and stored at -80°C. This procedure generated HS2ST of >95% purity. Proteolytic removal of the MBP affinity tag from HS2ST (after re-cloning with MBP and 3C protease sites into the plasmid pOPINM) led to rapid HS2ST denaturation, based on rapid precipitation, so for the procedures described in this paper the MBP affinity tag was left intact. For SDS-PAGE, proteins were denatured in Laemmli sample buffer, heated at 95°C for 5 min and then analysed by SDS-PAGE with 10% (v/v) polyacrylamide gels. Gels were stained and destained using a standard Coomassie Brilliant Blue protocol.

### DSF-based fluorescent assays

Thermal shift /stability assays (TSAs) were performed using a StepOnePlus Real-Time PCR machine (Life Technologies) using SYPRO-Orange dye (Emission max. 570 nm, Invitrogen), with thermal ramping between 20 - 95°C in 0.3°C step intervals per data point to induce denaturation in the presence or absence of test biochemicals or small molecule inhibitors, as previously described [45]. HS2ST was assayed at a final concentration of 5 μM in 50 mM Tris–HCl (pH 7.4) and 100 mM NaCl. Final DMSO concentration in the presence or absence of the indicated concentrations of ligand was no higher than 4% (v/v). Normalized data were processed using the Boltzmann equation to generate sigmoidal denaturation curves, and average *T*_m_/Δ*T*_m_ values were calculated as described using GraphPad Prism software [45].

### Microfluidics-based sulphation assay

N-sulphated, fluorescein-tagged hexasaccharide glycan substrate (GlcNS-GlcA-GlcNS-IdoA-GlcNS-GlcA-fluorescein, where S=sulphation) containing either L-IdoA or D-GlcA residues at the third residue from the reducing end (to which a linker and the fluorophore were conjugated) were both purchased from GLYCAN therapeutics (www.glycantherapeutics.com). The fluorescein group attached to the reducing end of the glycan substrate (GlcNS-GlcA-GlcNS-IdoA-GlcNS-GlcA-fluorescein, where S=sulphation) possesses a maximal emission absorbance of ~525 nm, which can be detected by the EZ Reader *via* LED-induced fluorescence. Heparin and heparan sulphate-derivatives were generated enzymatically or through direct chemical synthesis, as described previously [4, 46]. Non-radioactive microfluidic mobility shift carbohydrate sulphation assays were optimised in solution with a 12-sipper chip coated with SR8 reagent and a Perkin Elmer EZ Reader II system [47] using EDTA-based separation buffer and real-time kinetic evaluation of substrate sulphation. Pressure and voltage settings were adjusted manually to afford optimal separation of the sulphated product and non-sulphated hexasaccharide substrate, with a sample (sip) volume of 20 nl, and total assay times appropriate for the experiment. Individual sulphation assays were assembled in a 384 well plate in a volume of 80 µl in the presence of the indicated concentration of PAPS or various test compounds, 50 mM HEPES, 0.015 % (v/v) Brij-35 and 5 mM MgCl_2_ (unless specified otherwise). The degree of oligosaccharide sulphation was directly calculated using EZ Reader software by measuring the sulpho oligosaccharide:oligosaccharide ratio at each time-point. The activity of HS2ST enzymes in the presence of biochemicals and small molecule inhibitors was quantified in ‘kinetic mode’ by monitoring the amount of sulphated glycan generated over the assay time, relative to control assay with no additional inhibitor molecule (DMSO control). Data was normalized with respect to these control assays, with sulphate incorporation into the substrate limited to ~ 20 % to prevent depletion of PAPS and to ensure assay linearity. K*m* and IC_50_ values were determined by non-linear regression analysis with GraphPad Prism software.

### NMR-based oligosaccharide sulphation analysis

For NMR experiments, fluorescein-labelled hexasaccharide L-IdoA substrate and the HS2ST-catalysed sulphation product (10 µM) dissolved in 50 mM HEPES, pH 7.3, 5 mM MgCl_2_ and 0.002% (v/v) Brij-35 were lyophilised overnight and re-dissolved in an equivalent amount of D_2_O. NMR experiments were performed at 25°C on a Bruker Avance III 800 MHz spectrometers equipped with a TCI CryoProbe. 1D and 2D proton and TOCSY spectra (mixing time 80 ms) were measured using standard pulse sequences provided by the manufacturer. Spectra were processed and analysed using TopSpin 3.4 software (Bruker).

### HPLC-based oligosaccharide sulphation analysis

The fluorescein-labelled hexasaccharide L-IdoA substrate and the HS2ST-catalysed sulphation product (10 µM) were analyzed after anion exchange chromatography by HPLC as previously described (1). Oligosaccharides were digested in the presence of a mixture of heparitinase I, II and III. Samples were loaded on a Proteomix SAX-NP5 (SEPAX) column eluted with an NaCl gradient Column effluent was mixed (1:1) with 2% 2-cyanoacetamide in 250mM of NaOH and subsequently monitored with a fluorescence detector (JASCO; FP-1520) either at 346 nm excitation and 410 nm emission (detection of mono and disaccharides linked to cyanoacetamide) or at 490 nm excitation and 525 nm emission for detection of trisaccharides-linked to fluorescein.

### Small molecule screening assays

The PKIS chemical library (Supplementary Figure 6, designated as SB, GSK or GW compound sets) comprises 367 largely ATP-competitive kinase inhibitors, covering 31 chemotypes originally designed to inhibit 24 distinct protein kinase targets [48]. It was stored frozen as a 10 mM stock in DMSO. The library is characterised as highly drug-like (~70% with molecular weight <500 Da and clogP values <5). For initial screening, compounds dissolved in DMSO were pre-incubated with HS2ST for 10 minutes and then employed for DSF or sulphotransferase-based enzyme reactions, which were initiated by the addition of the universal sulphate donor PAPS. For inhibition assays, competition assays, or individual IC_50_ value determination, a compound range was prepared by serial dilution in DMSO, and added directly into the assay to the appropriate final concentration. All control experiments contained 4% (v/v) DMSO, which had essentially no effect on HS2ST activity. Individual chemicals and glycan derivatives were prepared and evaluated using NMR, HPLC, DSF or microfluidics-based assay protocols, as described above.

### Docking studies

Docking models for rottlerin, suramin and GW407323A were built using Spartan16 (https://www.wavefun.com) and energy minimised using the Merck molecular forcefield. GOLD 5.2 (CCDC Software;) was used to dock molecules [49], with the binding site defined as 10 Å around the 5’ phosphorous atom of PAP, using coordinates from chicken MBP-HS2ST PDB ID: 4NDZ [20]. A generic algorithm with ChemPLP as the fitness function [50] was used to generate 10 binding-modes per ligand in HS2ST. Protons were added to the protein. Default settings were retained for the “ligand flexibility” and “fitness and search options”, however “GA settings” was changed to 200%.

## RESULTS

### Analysis of human HS2ST ligand binding using a thermal stability assay (TSA)

To our knowledge, Differential Scanning Fluorimetry (DSF) has not previously been used to examine the thermal stability and shift profiles of sulphotransferases in the presence or absence of biochemical ligands, such as those related to the sulphate donor PAPS (Figure 1A). We purified recombinant HS2ST catalytic domain (amino acids 69 to 356) fused to an *N*-terminal Maltose Binding Protein (MBP) tag to near homogeneity (Figure 1B) and evaluated its thermal denaturation profile with the MBP tag still attached in the presence of PAPS, heparin or maltose (Figure 1C). As a control, we examined the profile of maltose-binding protein (MBP) incubated with the same chemicals (Figure 1D). Unfolding of MBP-HS2ST in buffer generated a biphasic profile, and the upper region of this profile could be positively shifted (stabilised) by incubation with the HS2ST co-factor PAPS or the known HS2ST-interacting oligosaccharide ligand heparin (Figure 1C). In contrast, maltose incubation with MBP-HS2ST induced the same characteristic stabilisation profile observed when MBP was incubated with maltose and then analysed by DSF (Figure 1D). As expected, neither PAPS nor heparin induced stabilisation of MBP, confirming that effects on MBP-HS2ST were due to interaction with the sulphotransferase domain, rather than the affinity tag of the recombinant protein (Figure 1D, relevant ∆T_m_ values presented in Figure 1E). Consistently, PAPS did not stabilise the catalytic domain of the ATP-dependent catalytic subunit of cAMP-dependent protein kinase (PKAc), which instead binds with high affinity to the co-factor Mg-ATP [45], inducing a ∆T_m_ value of >4°C (Figure 1F).

**Figure 1.**
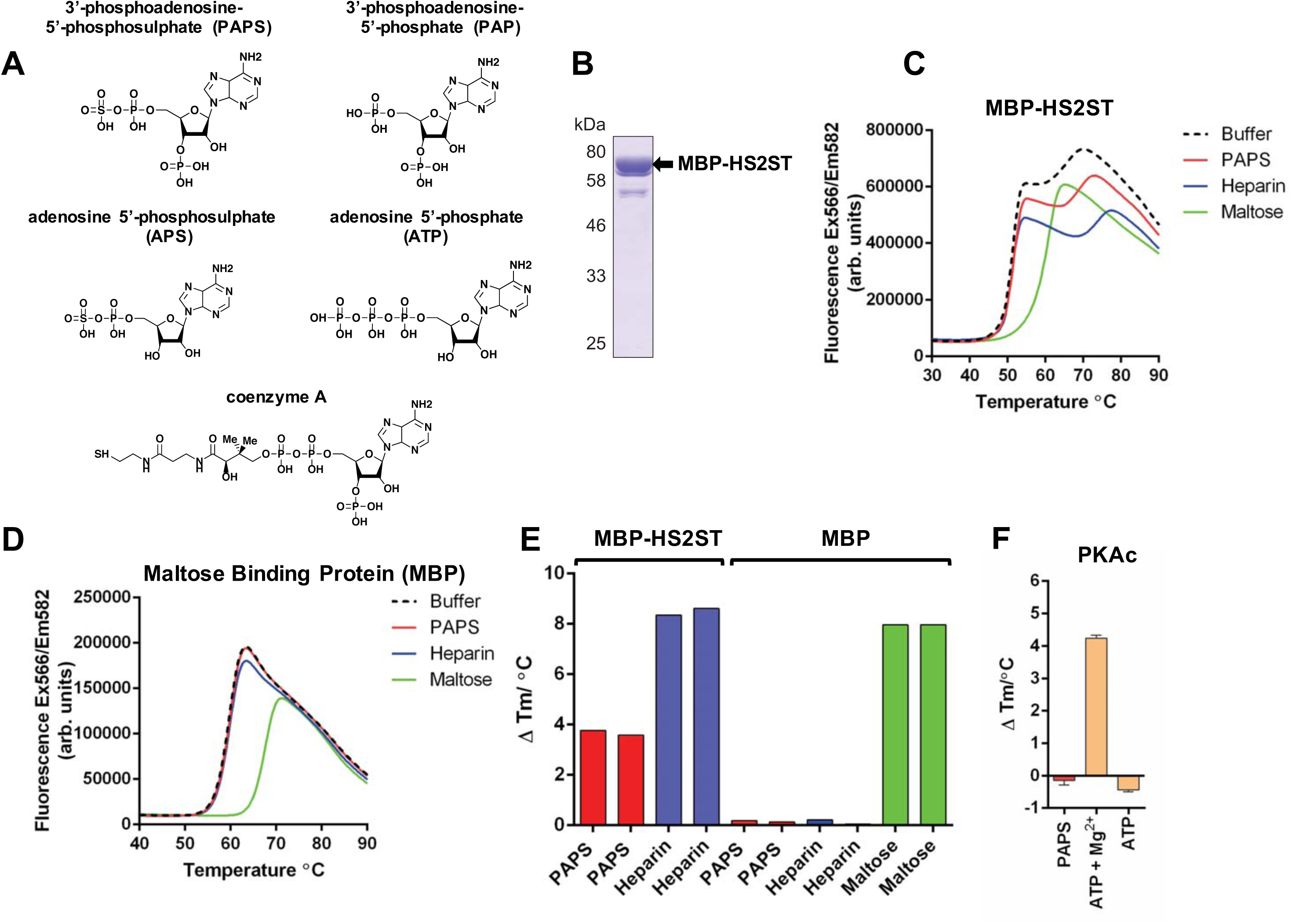
Analysis of purified recombinant MBP-HS2ST protein. **(A)** Structures of PAPS and PAPS-related biochemicals. **(B)** Coomassie blue staining of recombinant MBP-HS2ST1 protein. ~2 μg of purified enzyme was analysed after SDS-PAGE. **(C)** Thermal denaturation profiles of MBP-HS2ST (5 μM) and thermal shift in the presence of 0.5 mM PAPS (red), 10 μM heparin (blue) or 5 mM maltose (green). Buffer control is shown in black dashed lines. **(D)** Thermal denaturation profile of purified recombinant maltose binding protein (MBP). Experimental conditions as for (C). **(E)** T_m_ values measured for 5 μM MBP (squares) or MBP-HS2ST fusion protein (triangles) in the presence of 0.5 mM PAPS, 10 μM heparin or 5 mM maltose. ∆T_m_ values were obtained by DSF and calculated by subtracting control T_m_ values (buffer, no ligand) from the measured T_m_. **(F)** ∆T_m_ values relative to buffer addition for recombinant PKAc (5 μM) measured in the presence of 0.5 mM PAPS, 0.5 mM ATP or 0.5 mM ATP and 10 mM MgCl_2_. Similar results were seen in three independent experiments.

We next analysed the sensitivity of this assay for measuring HS2ST stability shifts over a wide range of PAPS concentrations, which confirmed dose-dependent stabilisation of recombinant HS2ST by PAPS, with detection of binding in the low micromolar range of the co-factor, equivalent to a molar ratio of ~1:1 HS2ST:PAPS (Supplementary Figure 1A). Subsequently, we explored the potential of this assay to detect binding of a putative IdoA-containing oligosaccharide substrate for HS2ST, confirming dose-dependent effects of this polymeric glycan over a range of concentrations, consistent with binding and conformational stability. Similar to PAPS, detection of binding was observed in the low micromolar range, equivalent to a molar ratio of ~1:1 HS2ST:glycan (Supplementary Figure 1B). We also evaluated binding of a panel of adenine-based cofactors (PAP and ATP), which suggested binding of divalent cation Mg^2+^ ions in an EDTA-sensitive manner (Supplementary Figure 1C), inducing a ΔTm of ~3°C, similar to that observed with the HS2ST co-factor PAPS. In contrast, removal of the sulpho moiety of PAPS, which creates the enzymatic end product PAP, was not deleterious to HS2ST binding (Supplementary Figure 2A), consistent with structural analysis of the enzyme [28]. Neither PAP nor PAPS binding required Mg^2+^ ions, although the effect on stabilisation with Mg^2+^ ions was additive (Supplementary Figures 1C and 2A). The non-functional enzyme co-factor APS, in which the 3’-phosphate group of adenine is absent, did not induce HS2ST stabilisation, confirming a requirement for this charged modification (Supplementary Figure 2A). We also established that CoA and acetyl CoA, which both contain a 3’-phosphoadenine moiety, clearly induced thermal stabilisation of HS2ST; loss of the 3’-phosphate group in dephospho CoA abolished this effect (Supplementary Figure 2A). Finally, we demonstrated that ATP, GTP and ADP, but not AMP or cAMP, were all effective at protecting HS2ST from thermal denaturation, suggesting that they are also HS2ST ligands (Supplementary Figure 2A).

### Analysis of human HS2ST glycan binding using TSA

To extend our HS2ST thermal analysis to identify potential glycan substrates, we evaluated enzyme stability in the presence of synthetic glycan chains of different lengths and sulphation patterns (Table 1). Of particular interest for further assay development, thermal shift (stabilisation) was detected in this assay when hexasaccharide (dp6) or a higher degree of polymerisation oligosaccharide was incubated with the enzyme (Supplementary Figure 2B), suggesting that a dp6 glycan might represent the shortest potential partner suitable for HS2ST binding, a prerequisite for enzymatic modification. Interestingly, highly sulphated hexameric glycans served as efficient HS2ST binding partners relative to the most highly sulphated heparin control, with a fully chemically sulphated I_2s,3s_A^6s^_3s_Ns hexamer inducing the highest HS2ST stability-shift amongst the chemically-modified derivatives assessed (Supplementary Figure 2C). Interestingly, a putative I_2OH_A^6OH^Ns substrate, which contains the 2-*O* moiety predicted to be the substrate for 2-O-sulphotransferases, also led to marked thermal stabilisation of HS2ST, suggestive of productive binding to HS2ST that permit it to be sulphated in the presence of PAPS (Supplementary Figure 2C).

### A novel microfluidic kinetic assay to directly measure oligosaccharide sulphation by HS2ST

In order to quantify the effects of various ligands on HS2ST enzyme activity, we sought to develop a new type of rapid non-radioactive solution assay that could discriminate the enzymatic incorporation of sulphate into a synthetic oligosaccharide substrate. Current protocols are time-consuming and cumbersome, requiring Mass Spectrometry, NMR or ^35^S-based radiolabelling/HPLC separation procedures. Importantly, we next tested whether a version of a I_2OH_A^6OH^NS hexasaccharide substrate coupled to a linker and fluorescein at the reducing end, which interacts with HS2ST (Supplementary Figure 2C), could also be enzymatically sulphated by HS2ST using ‘gold-standard’ NMR-based sulphation detection [44]. The fluorescent I_2OH_A^6OH^Ns could not be evaluated for binding to HS2ST by DSF, due to interference of the fluorescent group in the unfolding assay, which measures SYPRO-Orange fluorescence at a similar wavelength. Instead, to confirm sulphation of the fluorescein-labelled substrate, it was pre-incubated with PAPS and HS2ST to catalyse site-specific sulphation (Figure 2A). The NMR spectrum of the sulphated product compared to that of the non-modified substrate provided unequivocal evidence for sulphation at the 2-*O* position of the sugar, most notably due to the diagnostic shifts of anomeric H-1 and H-2 protons in the presence of the 2-*O*-sulphate group linkage to the carbon atom (Figure 2B and Supplementary Figure 3). The 2-*O* sulphated IdoA hexameric product was also confirmed using an established HPLC-based approach [51], which demonstrated stoichiometric sulphation of an enzyme-derived substrate derivative (Supplementary Figure 4).

**Figure 2.**
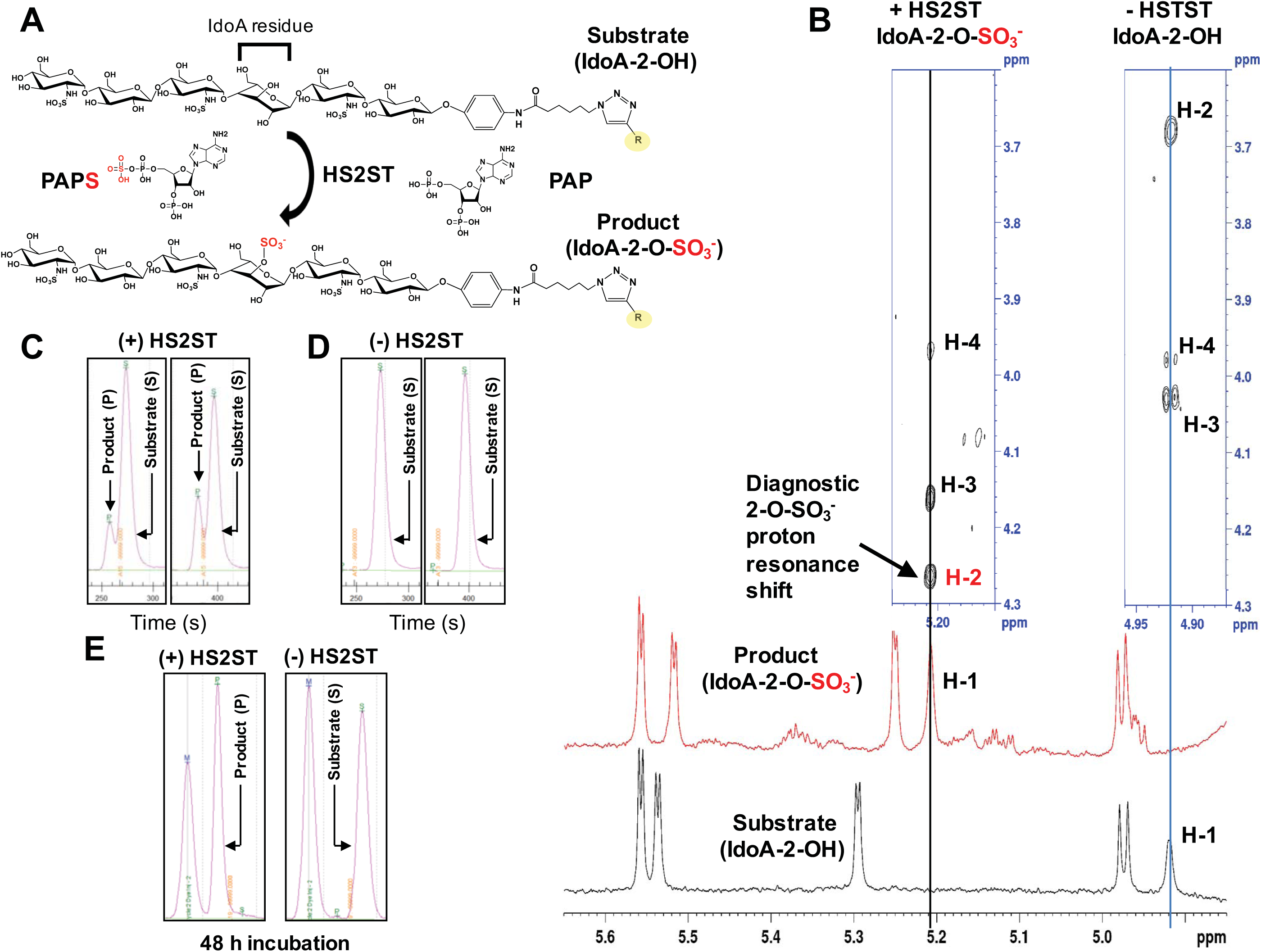
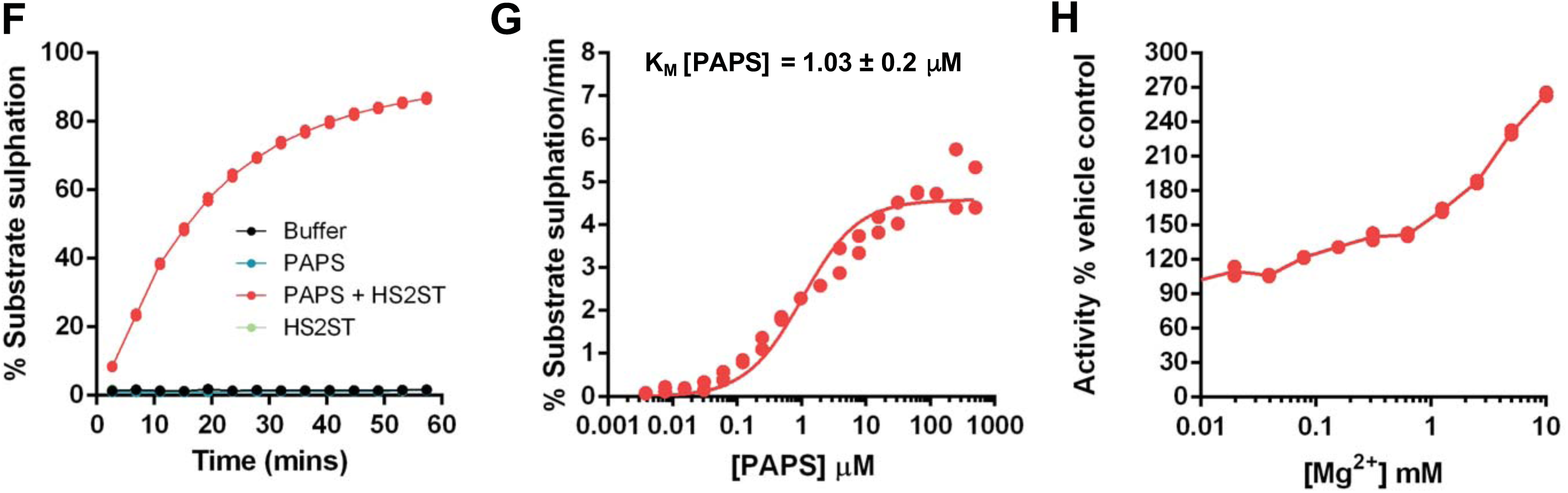
Development of a novel microfluidic mobility shift assay to quantify HS2ST enzymatic activity. **(A)** Schematic showing PAPS-dependent sulphate incorporation into the fluorescein-labelled hexasaccharide IdoA substrate by HS2ST, with the concomitant generation of PAP. R=fluorescein. **(B)** NMR analysis of the non-sulphated and sulphated hexasaccharides. The addition of a 2-O-sulphate group to the iduronate (L-IdoA) residue of the fluorescent hexasaccharide results in a significant chemical shift change, most notably to the anomeric proton (H-1) and that of H-2 attached to the sulphated carbon atom of L-IdoA, in agreement with expected values from the literature [44]. ^1^H NMR spectrum of non-sulphated substrate (bottom spectrum, black) and sulphated product (upper spectrum, red). Distinct L-IdoA protons (H-3 and H-4 of the spin system) were identified by TOCSY and are shown vertically above their respective H-1 signals (for the non-sulphated substrate, right blue boxed, and for the sulphated product, left blue boxed). The full carbohydrate proton spectra are shown in Supplementary Figure 3. **(C, D)** Screen shots of EZ reader II raw data files, demonstrating that HS2ST induces a rapid mobility change in the IdoA-containing fluorescent hexasaccharide. Separation of the higher mobility, sulphated (product, P) from the lower mobility (substrate, S) hexasaccharide occurs as a result of enzymatic substrate sulphation (left panels 180 s assay time, right panels 240 s assay time), as demonstrated by omission of HS2ST from the assay (-HS2ST). Assays were initially performed at 20°C using 90 nM of purified HS2ST, 2 μM fluorescein-labelled hexasaccharide substrate and 500 μM PAPS. **(E)** Stoichiometric sulphate-labelling of IdoA-containing fluorescein-labelled hexasaccharide. Reactions was performed with 0.6 μM HS2ST, 375 μM IdoA-hexasaccharide substrate and 1 mM PAPS and incubated at room temperature for 48 h. The reaction was spiked with an additional 0.5 mM (final concentration) of PAPS after 24 h of incubation. M = non-sulphated marker substrate. A final hexasaccharide concentration of 2 μM was analysed by fluorescent sulphation mobility assay. **(F)** Analysis of time-dependent sulphate incorporation into 2 μM IdoA-containing fluorescein-conjugated hexasaccharide. Percentage sulphation was calculated from the ratio of substrate hexasaccharide to product (2-*O*-sulpho)-hexasaccharide at the indicated time points in the presence or absence of 20 nM HS2ST and 10 µM PAPS. **(G)** Calculation of K*m* [PAPS] value for HS2ST. PAPS concentration was varied in the presence of a fixed concentration of HS2ST (20 nM), and the degree of substrate sulphation calculated from a differential kinetic analysis, n=2 assayed in duplicate. **(H)** Duplicate HS2ST assays conducted in the presence of increasing concentrations of activating Mg^2+^ ions. Activity is presented in duplicate relative to buffer controls. Similar results were seen in several independent experiments.

Next, we evaluated the incorporation of the sulphate moiety from PAPS into a fluorescently-labelled glycan substrate using a microfluidic assay that detects real-time changes in substrate covalent modification (notably the introduction of a negative charge) when an electric field is applied to the solution reaction. This ratiometric assay, which we and others have previously employed to detect the formal double negative charge induced by real-time peptide phosphorylation [42, 52–54], was able to detect real-time incorporation of sulphate into the oligosaccharide substrate, based on the different retention time of the product compared to the substrate (Figure 2C). No sulphated product was detected in the absence of HS2ST (Figure 2D), and prolonged incubation of substrate with HS2ST led to stoichiometric conversion of the substrate into the fully sulphated product (P), which migrated very differently to the substrate (S) ‘marker’ (Figure 2E). Analysis of product/(product + substrate) ratios of the peak heights allowed us to monitor sulphation over any appropriate assay time (Figures 2F), and the degree of sulphation could easily be varied as a function of PAPS concentration in the assay. Furthermore, no sulphated product was detected in the presence of buffer or PAPS alone (Figure 2F), allowing us to determine a K*m* value of ~1 µM for PAPS-mediated substrate hexasaccharide sulphation (Figure 2G). We also noted that high (>1 mM) concentrations of Mg^2+^ ions led to concentration-dependent increases in enzyme HS2ST activity (Figure 2H), consistent with the effects of Mg^2+^ ions identified in DSF assays (Supplementary Figure 2A). Finally, we confirmed that sulphation was optimal when an appropriate modifiable IdoA substrate was present, with sulphation reduced by >90% when a GlcA residue was incorporated into the substrate instead (compare Supplementary Figures 5A and 5B).

### Screening for small molecule inhibitors of HS2ST using DSF and microfluidic technology

The discovery of HS2ST inhibitors is hindered by a lack of a rapid and quantifiable assay for the facile detection of sulphate modification using a close mimic of a physiological substrate. Our discovery that a synthetic HS2ST glycan substrate could be readily sulphated and detected by enzymatic assay in solution, without the need for HPLC, NMR or radioactive procedures, meant that this approach might now be optimised for the discovery of small molecule HS2ST inhibitors. We first evaluated the ability of an unlabelled (non-fluorescent) hexameric glycan substrate that lacked sulphate at the 2-*O* position, or a non-substrate that was fully sulphated at all potential acceptor sites, to act as HS2ST inhibitors in our fluorescent glycan sulphation assay. As detailed in Figure 3A, the fully sulphated glycan was an extremely potent inhibitor, interfering with HS2ST-dependent sulphation of the substrate with an IC_50_ value of <10 nM, consistent with tight binding to the enzyme, as previously established using DSF (Supplementary Figure 2C). In contrast, a less highly sulphated substrate was still able to compete with the fluorescent substrate in a dose-dependent manner (fixed at 2 µM in this assay), as indicated by the IC_50_ value of <100 nM. We next compared the effects of PAP, ATP, CoA and dephospho-CoA, which all exhibit thermal stabilisation of HS2ST in DSF assays (Supplementary Figure 2A). Interestingly, PAP (IC_50_ ~2 µM), CoA (IC_50_ = 65 µM) and ATP (IC_50_ = 466 µM) were HS2ST inhibitors, whereas dephospho CoA (which lacks the 3’-phosphate moiety in CoA) was not (Figure 3B). Increasing the concentration of PAPS in the assay led to a decrease in the level of inhibition by both PAP and CoA (Figure 3C), suggesting a PAPS-competitive mode of inhibition, as predicted from the various shared chemical features of these molecules (Figure 1A).

**Figure 3.**
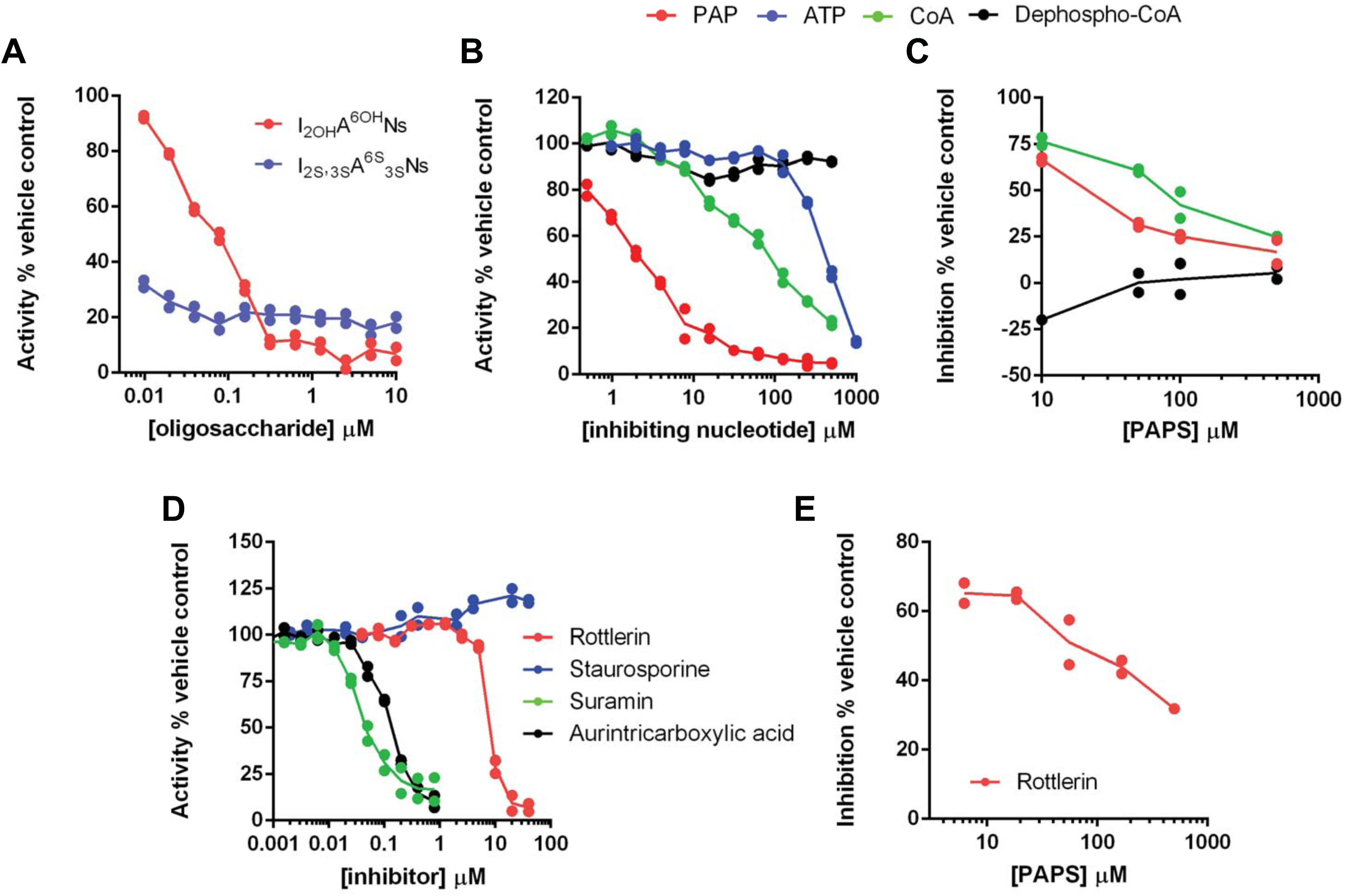
Microfluidic sulphotransferase assay to measure inhibition of HS2ST activity *in vitro.* Assays were performed using 20 nM HS2ST and the extent of substrate sulphation was determined after 15 mins incubation at room temperature as described in the legend to Figure 2. Dose-response curves for inhibition of HS2ST activity by **(A)** modified heparin derivatives containing different sulphation patterns (assayed in the presence of 0.5 mM MgCl_2_) or **(B)** nucleotides (assayed in the absence of MgCl_2_). Assays contained HS2ST and 10 μM PAPS and the indicated concentration of inhibitory ligand or buffer. **(C)** Inhibition of HS2ST activity by fixed 10 μM PAP, 0.5 mM CoA or 0.5 mM dephospho-CoA in the presence of increasing concentration of PAPS. Inhibition is calculated as a function of no inhibitor for each concentration of PAPS in the absence of MgCl_2_. **(D)** Evaluation of small molecule HS2ST inhibitory profiles in the presence of 10 μM PAPS. **(E)** Inhibition HS2ST activity by 20 μM rottlerin in the presence of varied concentrations of PAPS, suggesting a competitive mode of inhibition. Similar results were seen in multiple experiments.

Recent studies have demonstrated that PAPS-dependent tyrosyltransferases are inhibited by several non-nucleotide-based polyanionic chemicals [55]. However, to our knowledge, the inhibition of carbohydrate sulphotransferases by such compounds has not been reported. Using our microfluidic assay, we confirmed that the polysulphated compound suramin (an inhibitor of angiogenesis) and the polyaromatic polyanion aurintricarboxylate (an inhibitor of protein:nucleic acid interactions, DNA polymerase and topoisomerase II) demonstrated nanomolar inhibition of HS2ST, with IC_50_ values of 40 ± 1 nM and 123 ± 7 nM respectively (Figure 3D). In addition, the non-specific protein kinase inhibitor rottlerin also inhibited HS2ST with an IC_50_ of 6.4 µM. Increasing the concentration of PAPS in the sulphation assay decreased the inhibitory effect, consistent with a competitive mode of HS2ST inhibition for rottlerin (Figure 3E).

### Protein kinase inhibitors are a new class of potential broad-spectrum HS2ST inhibitor

The finding that the non-specific kinase inhibitor rottlerin [56] was a micromolar inhibitor of HS2ST was of particular interest, especially given the remarkable progress in the development of kinase inhibitors as chemical probes, tool compounds and, latterly, clinically-approved drugs. Similarities between ATP and PAPS (Figure 1A), and the finding that ATP can both bind to, and inhibit, HS2ST activity (Supplementary Figure 2A and Figure 3B) raised the possibility that other ATP-competitive protein kinase inhibitors might also interact with HS2ST. In order to exploit our screening capabilities further, we established a 384-well assay to evaluate inhibition of PAPS-dependent glycan sulphation by HS2ST. The Published Kinase Inhibitor Set (PKIS) is a well-annotated collection of 367 high-quality ATP-competitive kinase inhibitor compounds that are ideal for compound repurposing or the discovery of new chemical ligands for orphan targets. We screened PKIS using DSF and enzyme-based readouts (Figures 4A and B respectively). As shown in Figure 4A, when screened at 40 µM compound in the presence of 5 µM HS2ST, only a small percentage of compounds induced HS2ST stabilisation or destabilisation at levels similar to that seen with an ATP control. We focussed on compounds inducing HS2ST ∆T_m_ values between + 0.5°C and - 0.5°C, and re-screened each ‘hit’ compound using ratiometric HS2ST enzyme assays at a final compound concentration of 40 µM. We reported the enzyme activity remaining compared to DMSO, with rottlerin (IC_50_ = ~8 µM), suramin (IC_50_ = ~20 nM) and aurintricarboxylate (IC_50_ = ~90 nM) as positive controls (Figure 4B and Supplementary Figures 6 and 7). We also included the compound GW406108X in our enzyme assay, since it was structurally related to several ‘hit’ compounds from the DSF screen. As shown in Figure 4C, the three PKIS compounds with the highest inhibitory activity (red) exhibited IC_50_ values of between 20-30 µM towards HS2ST in the presence of 1 μM PAPS, similar to inhibiton by rottlerin. Of particular interest, these three compounds were amongst the top ~10% of compounds in terms of their ∆T_m_ values (red spheres, Figure 4A). Chemical deconvolution of compounds revealed that all three were closely-related members of a class of oxindole-based RAF protein kinase inhibitor (Figure 4A). Subsequently, one other related indole RAF inhibitory compound from PKIS, GW305074, was also shown to be a mid-micromolar HS2ST inhibitor, whereas the related oxindole GW405841X (Supplementary Figure 7) did not inhibit HS2ST at any concentration tested. Finally, we used combined DSF and enzyme assays to evaluate a broader panel of kinase inhibitors (Supplementary Figure 8). However, neither the pan-kinase inhibitor staurosporine, nor several FDA-approved tyrosine kinase inhibitors bound HS2ST at any concentration tested. Moreover, chemically diverse RAF inhibitors, including clinical RAF compounds such as dabrafenib and vemurafenib were unable to inhibit HS2ST *in vitro*, even at concentrations as high as 400 µM in our sensitive HS2ST enzyme assay (Supplementary Figure 8B).

**Figure 4.**
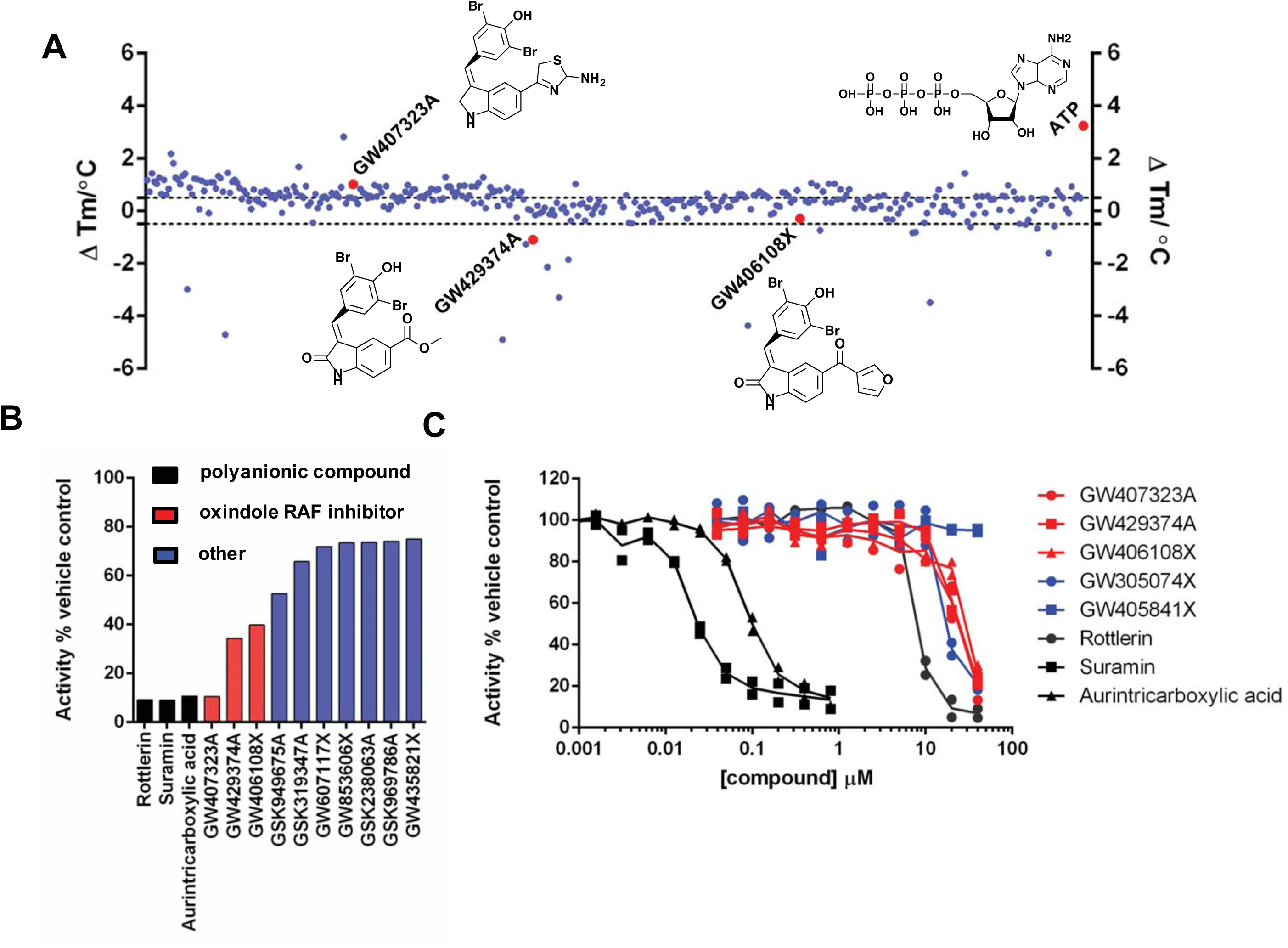
Mining the PKIS inhibitor library for HS2ST inhibitor compounds. **(A)** Evaluation of small molecule ligands in a high-throughput HS2ST DSF assay. 5 μM HS2ST was screened in the presence or absence of 20 μM compound. The final concentration of DMSO in the assay was 4 % (v/v). ∆T_m_ values (positive and negative) were calculated by subtracting the control T_m_ value (DMSO alone) from the measured T_m_ value. Data shown on a scatter plot of the mean ∆T_m_ values from two independent DSF assays. **(B)** Enzymatic analysis of HS2ST inhibition by selected PKIS compounds. HS2ST (20 nM) was incubated with the indicated PKIS compound (40 μM) in the presence of 10 μM PAPS for 15 mins at room temperature. HS2ST sulphotransferase activity was assayed using the fluorescent hexasaccharide substrate and normalised to DMSO control (4 % v/v). **(C)** Full dose-response curves for selected compounds. HS2ST (20 nM) was incubated with increasing concentration of inhibitor in the presence of 1 μM PAPS for 15 mins at 20°C. HS2ST activity calculated as above. Data from two independent experiments are combined. Similar results were seen in two independent experiments.

### Docking analysis of HS2ST ligands

The X-Ray structure (PDB ID:4NDZ) of trimeric chicken MBP-HS2ST fusion protein bound to non-sulphated PAP (Adenosine-3’-5’-diphosphate, a potent HS2ST inhibitor identified in this study) and a polymeric oligosaccharide, have previously been reported [20, 28], and we employed a 3.45 Å structural dataset to dock rottlerin, suramin and the most potent oxindole ‘hit’ (GW407323A, Figure 4B) from the screen, into the extended enzyme active site. As shown in Figure 5A, HS2ST possesses substrate-binding features that accommodates an extended oligosaccharide that place it in close proximity to the desulphated PAP end-product, which substitutes for the endogenous PAPS co-factor during crystallisation. The 3’-phosphoadenine moiety of PAP also helps anchor the nucleotide in an appropriate position. A molecular docking protocol for PAP in HS2ST was developed that matched the crystallographic binding pose of PAP extremely well (RMSD 0.31 Å, Figure 5B). By comparing a crystallised ligands (ADP) with docked rottlerin, suramin and GW407323A, we confirmed that compounds could be docked into the active site of HS2ST broadly corresponding to either the PAPS-binding region (rottlerin and GW407323A, Figures 5C and D) or bridging both the substrate and co-factor binding sites (suramin, Figure 5E). In these binding modes, compounds make a number of stabilising interactions that permit them to compete with PAPS or oligosaccharide substrate for binding to HS2ST (Figure 5C). For example, rottlerin is predicted to form a hydrogen bond with the amide backbone of Thr1290, GW407323A has multiple potential hydrogen bonding interactions with residues including Arg1080, Asn1112 and Ser1172, whilst suramin is predicted to form hydrogen bonds with residues Asn1091, Tyr1094 and Arg1288, targeting this highly elongated inhibitor to both active sites.

**Figure 5.**
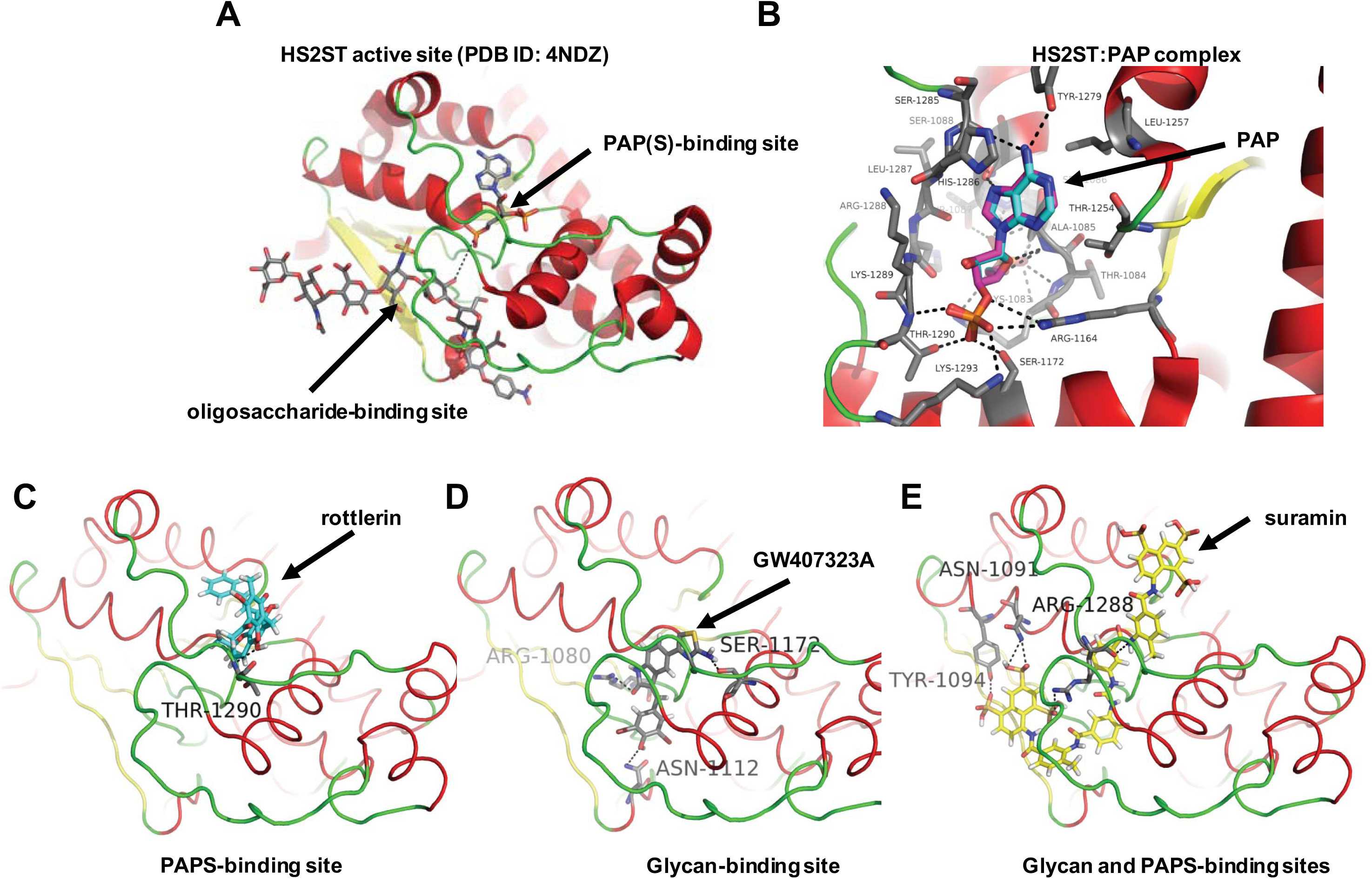
Molecular docking analysis of HS2ST with small molecule inhibitor compounds. **(A)** Structural representation of the catalytic domain of chicken MBP-HS2ST crystallised with bound heptasaccharide and non-sulphated PAP co-factor (Protein rendered as a cartoon. Red – α helix, yellow – β sheet, green – loop. PAP (Adenosine-3’-5’-diphosphate) and heptasaccharide are rendered as coloured sticks. Grey – carbon, red, oxygen, blue – nitrogen, yellow – sulphur. Black dotted line indicates close proximity of glycan 2-OH group and PAP. **(B)** Structure of HS2ST with near identical crystallographic (carbons in cyan) and docking (carbons in purple) poses of PAP (Protein rendered as a cartoon. Red – α helix, yellow – β sheet, green – loop. PAP rendered as coloured sticks. Cyan/Grey/Purple – carbon, red, oxygen, blue – nitrogen, dark yellow – sulphur). Black dotted lines indicate hydrogen bonds. Molecular Docking of **(C)** rottlerin, **(D)** the indole RAF inhibitor GW407323A or **(E)** suramin into the HS2ST catalytic domain (Protein depicted as a cartoon. Red – α helix, yellow – β sheet, green – loop. Docked molecules coloured as sticks. Pink/Yellow/Salmon/Grey – carbon, red, oxygen, blue – nitrogen, dark yellow – sulphur, white – hydrogen). Black dotted lines indicate hydrogen bonds).

## DISCUSSION

In this paper, we report a simple method for the detection of enzyme-catalysed glycan sulphation using a model IdoA-containing hexasaccharide fused to a reducing-end fluorophore. The chemical similarity between ATP, a universal phosphate donor, and PAPS, a universal sulphate donor, led us to investigate whether enzymatic glycan sulphation could be detected using a high-throughput kinetic procedure previously validated for peptide phosphorylation by ATP-dependent protein kinases. We focussed our attention on HS2ST, which transfers sulphate from PAPS to the 2-*O* position of IdoA during heparan sulphate biogenesis in the secretory pathway.

To facilitate rapid purification of recombinant HS2ST, the enzyme was expressed as an N-terminal MBP fusion protein, and we confirmed that it was folded, and could bind to a variety of exogenous ligands including PAPS and PAP, the end product of the sulphotransferase reaction. Protein kinases are also known to bind to their end product (ADP), and kinase structural analysis has long taken advantage of the stability of kinase and ADP complexes for protein co-crystallisation. Similar co crystallisation approaches revealed the structure of HS2ST, and related sulphotransferases, in complex with PAP and model saccharide substrates [20, 21], and our study extends these approaches, by revealing a competitive mode of HS2ST interaction with a variety of 3’-phosphoadenosine-containing nucleotides, including Coenzyme A (CoA). They also suggest that generalised docking of a 3’phosphoadenosine moiety is a feature of HS2ST that could be mimicked using other small molecule inhibitors. DSF-based thermal shift assays are ideal for the analysis of a variety of proteins and ligands, including growth factors [4, 57], protein kinase domains [45, 54, 58], pseudokinase domains [59], BH3 [60] and bromodomain-containing proteins [61]. However, to our knowledge, this is the first report to demonstrate the utility of a DSF-based strategy for the analysis of a sulphotransferase.

### Competitive HS2ST inhibition by biochemical ligands

By developing a new type of rapid, kinetic glycan sulphation assay, we confirmed that many HS2ST ligands also act as competitive inhibitors of PAPS-dependent oligosaccharide sulphation, setting the stage for a broader screening approach for the discovery of HS2ST inhibitors. Standard assays for carbohydrate sulphation utilise HPLC-based detection of ^35^S-based substrate sulphation derived from ^35^S-lablled PAPS, requiring enzymatic co-factor synthesis and time-consuming radioactive solid-phase chromatography procedures [20, 35, 41]. Whilst enzymatic deconvolution, MS and NMR-based procedures remain useful for mapping sulphation patterns in complex (sometimes unknown) glycan polymers, these procedures are very time-consuming and relatively expensive. In contrast, our finding that sulphation can be detected using a simple glycan mobility shift assay, and then quantified in real time by comparing the ratio of a sulphated and non-sulphated substrate, is rapid, reproducible and highly cost effective. Our kinetic assay makes use of a commercial platform originally developed for the analysis of peptide phosphorylation or peptide proteolysis, which allows for the inclusion of high concentrations of non-radioactive co-factors, substrates and ligands in assays [58]. Consequently, we were able to use this technology to derive a K*m* value for PAPS in our standard HS2ST assay of 1.0 µM (Figure 2G), slightly lower than the reported literature value of 18.5 µM for HS2ST using desulphated heparin as substrate [20], but similar to the reported literature value of ~4.3 µM for the PAPS-dependent GlcNAc-6-sulphotransferase NodH from *Rhizobium melitoli* [35] and 1.5 and 10 µM for human hormone iodothyrosine sulphotransferases and tissue-purified tyrosyl sulphotransferase [62, 63]. In the course of our studies, we developed several new reagents, including a hexameric fluorescent substrate in which the IdoA residue was replaced by a GlcA residue (Supplementary Figure 4). Interestingly, a decreased rate of substrate modification was observed using this oligosaccharide substrate, consistent with the ability of HS2ST to sulphate either IdoA or GlcA [19], but with a preference for the former. Previous HPLC-based studies identified an N-sulpho group in the oligosaccharide substrate as a pre-requisite for catalysis, with subsequent preferential transfer of sulphate to the 2-*O* position of IdoA [20, 22, 28, 64]; these published observations are entirely consistent with our findings using a hexameric fluorescent substrate.

In the future, it might be possible to quantify other site-specific covalent modifications in complex glycans using fluorescent oligosaccharides that contain distinct sugar residues, and by employing mobility-dependent detection in the presence of a variety of enzymes. These could include 3-*O* and 6-*O* sulphotransferases [21] or structurally distinct glycan phosphotransferases, such as the protein-O-mannose kinase POMK/Sgk196 [65], which catalyses an essential phosphorylation step during biosynthesis of an α-dystroglycan substrate [66]. Using this general approach, the screening and comparative analysis of small molecule inhibitors of these distinct enzyme classes would be simplified considerably relative to current procedures.

### HS2ST inhibition by previously known kinase inhibitors, including a family of RAF inhibitor

Our finding that HS2ST was inhibited at sub-micromolar concentrations by the compounds suramin [67] and the DNA polymerase inhibitor aurintricarboxylic acid [68] was intriguing, and consistent with recent reports demonstrating inhibitory activity of these compounds towards tyrosyl protein sulphotransferases, which employ PAPS as a co-factor, but instead sulphate tyrosine residues in proteins [55]. During the course of our studies screening a panel of kinase inhibitors, we found that the non-specific kinase compound rottlerin is a low micromolar inhibitor of HS2ST *in vitro*, with inhibition dependent upon the concentration of PAPS in the assay, suggesting a competitive mode of interaction. Rottlerin (also known as mallotoxin) is a polyphenolic compound from *Mallotus philippensis*, and although originally identified as an inhibitor of PKC isozymes [69], possesses a wide variety of biological effects likely due to its non-specific inhibition of multiple protein kinases [56]. This lack of specificity prevents exploitation of rottlerin in cells as a specific probe, although our finding that HS2ST is a target of this compound opens up the possibility that this, or other, protein kinase inhibitors might also possess inhibitory activity towards HS2ST, either due to an ability to target the PAPS or oligosaccharide-binding sites in the enzyme. To evaluate these possibilities further, we screened PKIS, a collection of drug-like molecules with broad inhibitory activity towards multiple protein kinases. Interestingly, only 3 compounds (<1% of the library) consistently showed marked inhibitory activity at 40 µM in our HS2ST enzyme assay (Figure 4A, B and C, red). Remarkably, all three compounds belonged to the same benzylidene-1H-inol-2-one (oxindole) chemical class, which were originally reported as potent ATP-dependent RAF kinase inhibitors that block the MAPK signalling pathway in cultured cells [70]. Retrospectively, of all the related chemotypes present in the PKIS library, we confirmed that GW305074X (but not GW405841X) was also a low micromolar HS2ST inhibitor, consistent with the broad sensitivity of HS2ST to this optimised class of RAF inhibitor.

Although limited Structure Activity Relationships can be derived from our initial studies, these findings demonstrate that HS2ST inhibitors can be discovered, and that several of these inhibitors could be of broad interest to the sulphotransferase (and protein kinase) fields. Our study also validates previous observations from the turn of the century, in which carbohydrate inhibitors of NoDH sulphotransferase were reported from a low diversity kinase-directed library [35]. Surprisingly, this early breakthrough did not lead to the development of any glycan sulphotransferase tool compounds for cell-based analysis. However, our discovery that oxindole-based RAF inhibitors are also HS2ST inhibitors could provide new impetus for the design and synthesis of much more specific and potent HS2ST inhibitors from this class of RAF kinase inhibitor. A requirement for rapid progress during this process will be structure-based analysis of HS2ST in the presence of compounds, in order to determine mechanism and mode(s) of interaction. Our initial docking studies suggest similar binding modes for both rottlerin and the oxindole-based ligand GW407323A (Figure 5), with the potential for cross-over between PAPS and substrate-binding sites present on the surface of HS2ST. It will be intriguing to explore these binding modes by structural analysis and guided mutational approaches [71], to evaluate potential drug-binding site residues in HS2ST and tease-apart requirements for enzyme inhibition. It will also be important to evaluate whether compounds identified as *in vitro* HS2ST inhibitors, including RAF inhibitors, can also interfere with HS sulphation and downstream signalling in cells. Interestingly, suramin is a potent anti-angiogenic compound, and is reported to have cellular effects on FGF signalling [72], whereas aurintricarboxylate has multiple cellular effects currently attributed to nucleotide-dependent processes; attempting to link some of these cellular phenotypes to the inhibition of glycan sulphation is also a worthy experimental strategy.

## CONCLUSION

Our work raises the possibility that HS2ST inhibitors could be strategically developed following the successful blueprint laid down for protein kinase inhibitors in the previous decades. Dozens of sulphotransferases are found in vertebrate genomes, and the development of chemical biology approaches to rapidly inactivate Golgi membrane-bound sulphotransferases and induce targeted inhibition of sulphation has been stymied by a lack of tool compounds, which have the opportunity to revolutionise cell biology when properly validated [73, 74]. We propose that if such compounds can be developed, perhaps by the discovery of new inhibitors, or through chemical manipulation of the leads reported in this study, then a new era in sulphation-based cell biology might be on the horizon. By generating tools to chemically control glycan sulphation modulated by HS2ST directly, inhibitor-based interrogation of sulphation-dependent enzymes could also have significant impact in many active areas of translational research.

## ACKNOWLEDGEMENTS

This work was funded by a BBSRC Tools and Resources Development Grant (BB/N021703/1) and a Royal Society Research Grant (to PAE), a European Commission FET-OPEN grant (ArrestAD no.737390) to DPG, SC, DGF and PAE, North West Cancer Research (NWCR) grants CR1088 and CR1097 and a NWCR endowment (to DGF). VP is supported by NIH Small Business Innovation Research Contract HHSN261201500019C. The SGC is a registered charity (number 1097737) that receives funds from AbbVie, Bayer Pharma AG, Boehringer Ingelheim, Canada Foundation for Innovation, Eshelman Institute for Innovation, Genome Canada, Innovative Medicines Initiative (EU/EFPIA) [ULTRA-DD grant no. 115766], Janssen, Merck KGaA Darmstadt Germany, MSD, Novartis Pharma AG, Ontario Ministry of Economic Development and Innovation, Pfizer, São Paulo Research Foundation-FAPESP, Takeda, and The Wellcome Trust [106169/ZZ14/Z].

## AUTHOR CONTRIBUTIONS

PAE obtained BBSRC grant funding with DGF and EAY. PAE, DPB, EAY, ILB, CEE, DPG, SC and NGB designed and executed the experiments. VP, JL, CW, DHD and WJZ provided critical reagents, compound libraries, protocols and critical advice. PAE wrote the paper with contributions and final approval from all of the co-authors.

---

**Supplementary Figure 1.**
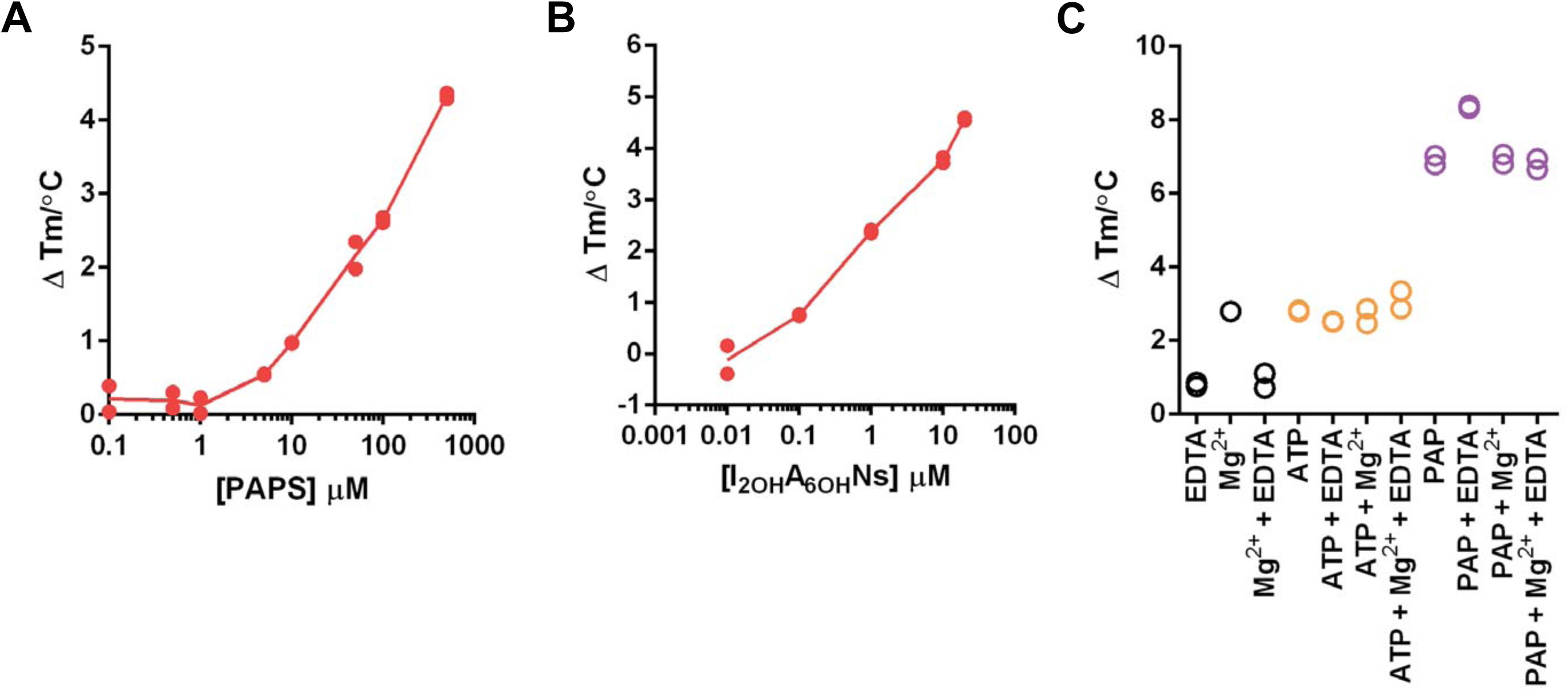
Thermal stability analysis of MBP-HS2ST. Concentration-dependent thermal profiling of MBP-HS2ST in the presence of **(A)** PAPS and the chemically-modified heparin derivative I_2OH_A^60H^N_S_ (compound 7, see Table 1). **(B)** TSA of 5 μM MBP-HS2ST measured in the presence of the indicated concentration of PAPS or I_2OH_A^60H^N_S_. ∆T_m_ values were calculated by DSF as previously described. **(C)** TSA assay showing changes in MBP-HS2ST thermostability induced by PAP and ATP, and the effects of EDTA and Mg^2+^. Thermal stability of HS2ST was measured as a function of compound binding by DSF. ∆T_m_ values of HS2ST protein (5 μM) incubated with 0.5 mM of the indicated nucleotide ± 10 mM MgCl_2_ ± 10 mM EDTA are shown. (Describe Colours)

**Supplementary Figure 2.**
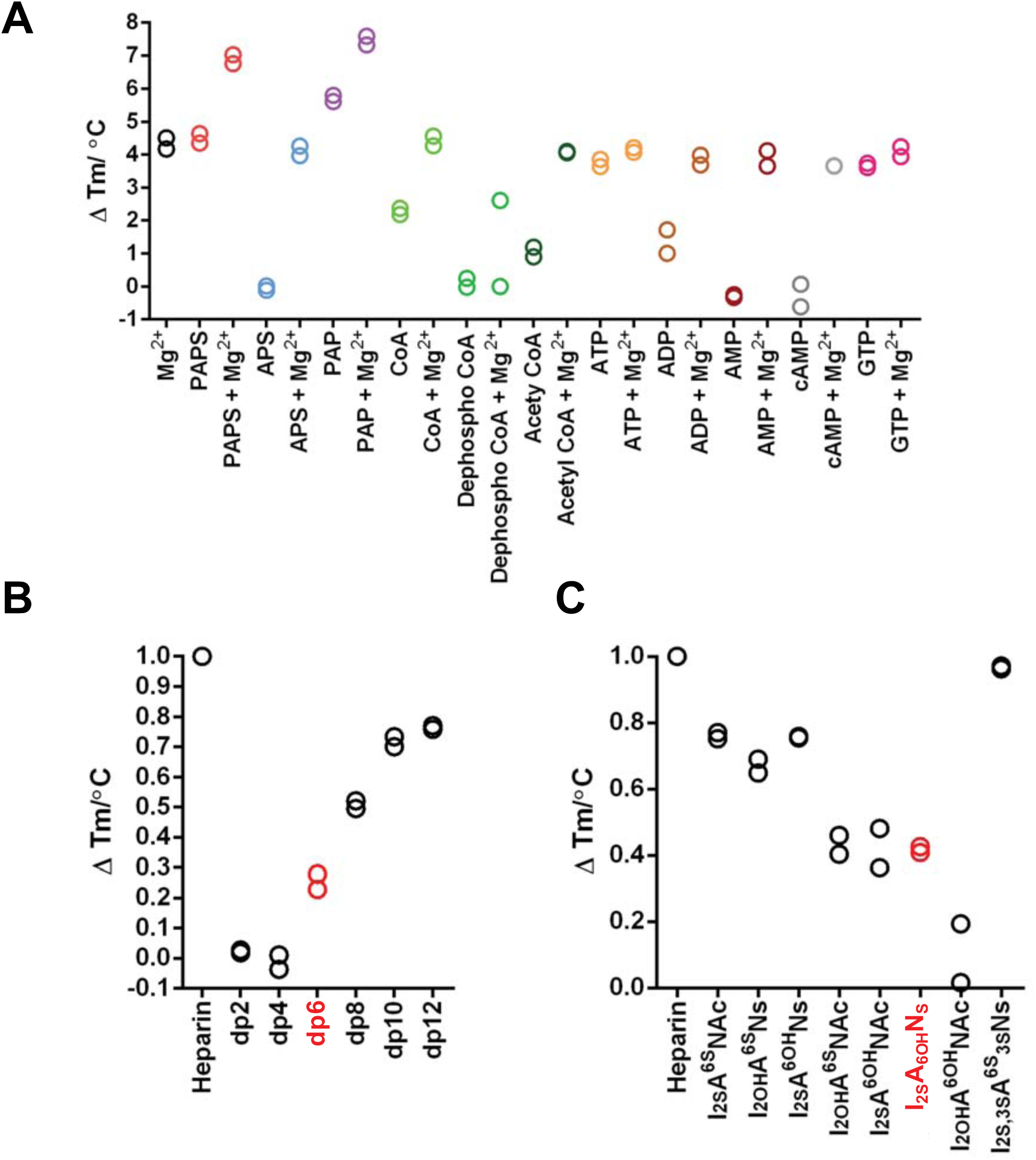
MBP-HS2ST Nucleotide and polysaccharide analysis. **(A)** TSA showing MBP-HS2ST binding of nucleotides by DSF. Thermal stability was measured as a function of nucleotide binding by DSF. ∆T_m_ values of HS2ST protein (5 μM) incubated with 0.5 mM of the indicated nucleotide ± 10 mM MgCl_2_ are shown. DSF analysis showing thermal shift (stabilization) of 5 μM HS2ST in the presence of 10 μM size separated oligosachharide fragments, dp (degree of polymerisation) equivalent to disacchardide (dp2), tetrasaccharide (dp4), hexasaccharide (dp6), octasaccharide (dp8), decasaccharide (dp10) or dodecasaccharide (dp12) **(B)** or chemically-modified heparin derivatives **(C)**. The minimal hexasaccharide binding substrate in (B) and the putative HS2ST substrate I_2OH_A^6OH^Ns in (C) are both shown in red. ∆T_m_ values (calculated as previously described) are normalized relative to heparin. dp=degree of polymerisation.

**Supplementary Figure 3.**
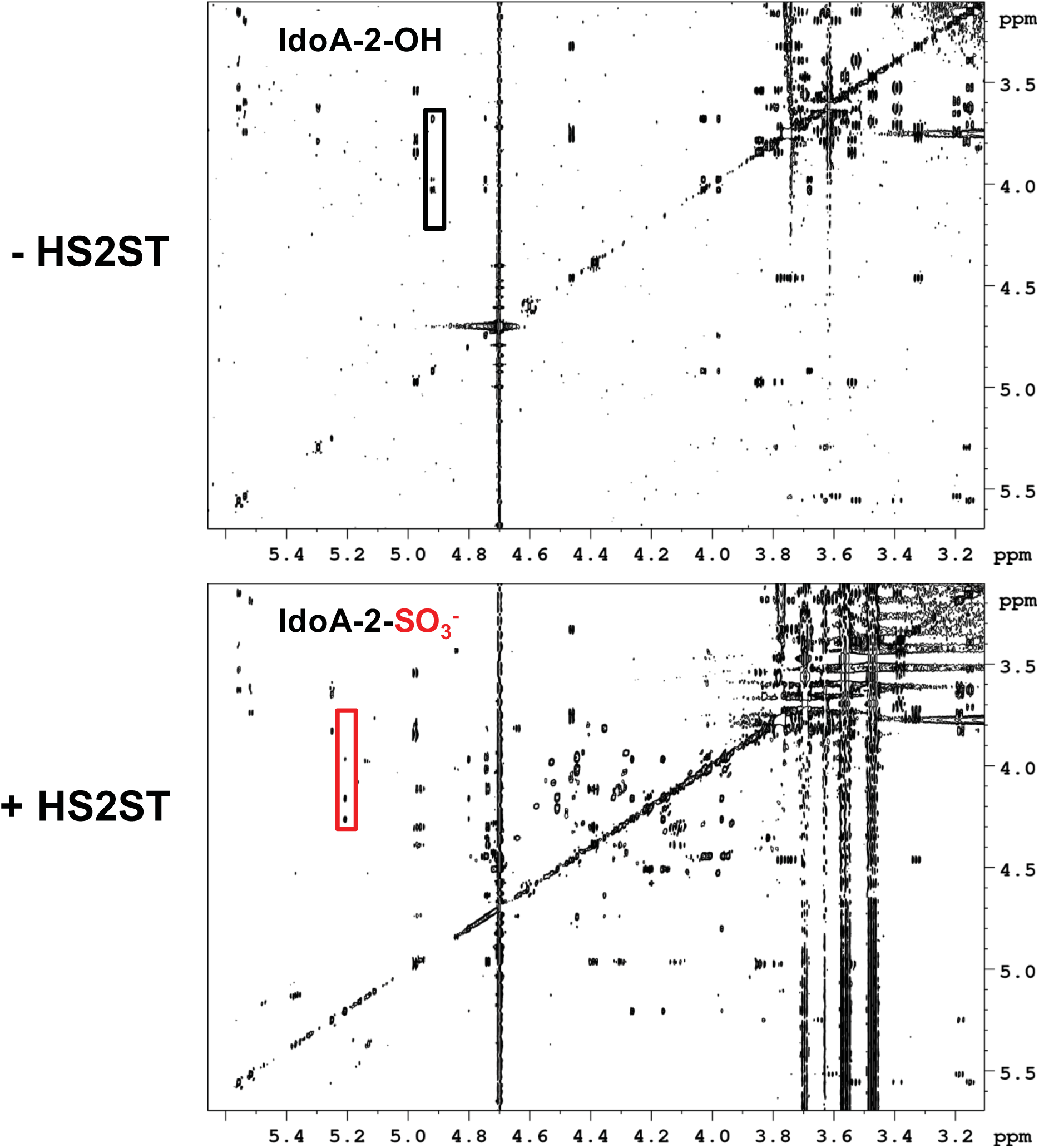
NMR spectra of sulphated and non-sulphated fluorescent polysaccharide substrate. TOCSY spectra of the L-IdoA-containing hexameric fluorescein-labelled HS2ST substrate (top) and the 2-O-sulphated product (bottom) generated by incubation with HS2ST, including the full spectrum of all carbohydrate hydrogens detected. Selected spectral regions, including the diagnostic shift caused by 2-O-sulphation, are expanded in Figure 2B in the main text, and are highlighted here by black and red boxes respectively.

**Supplementary Figure 4.**
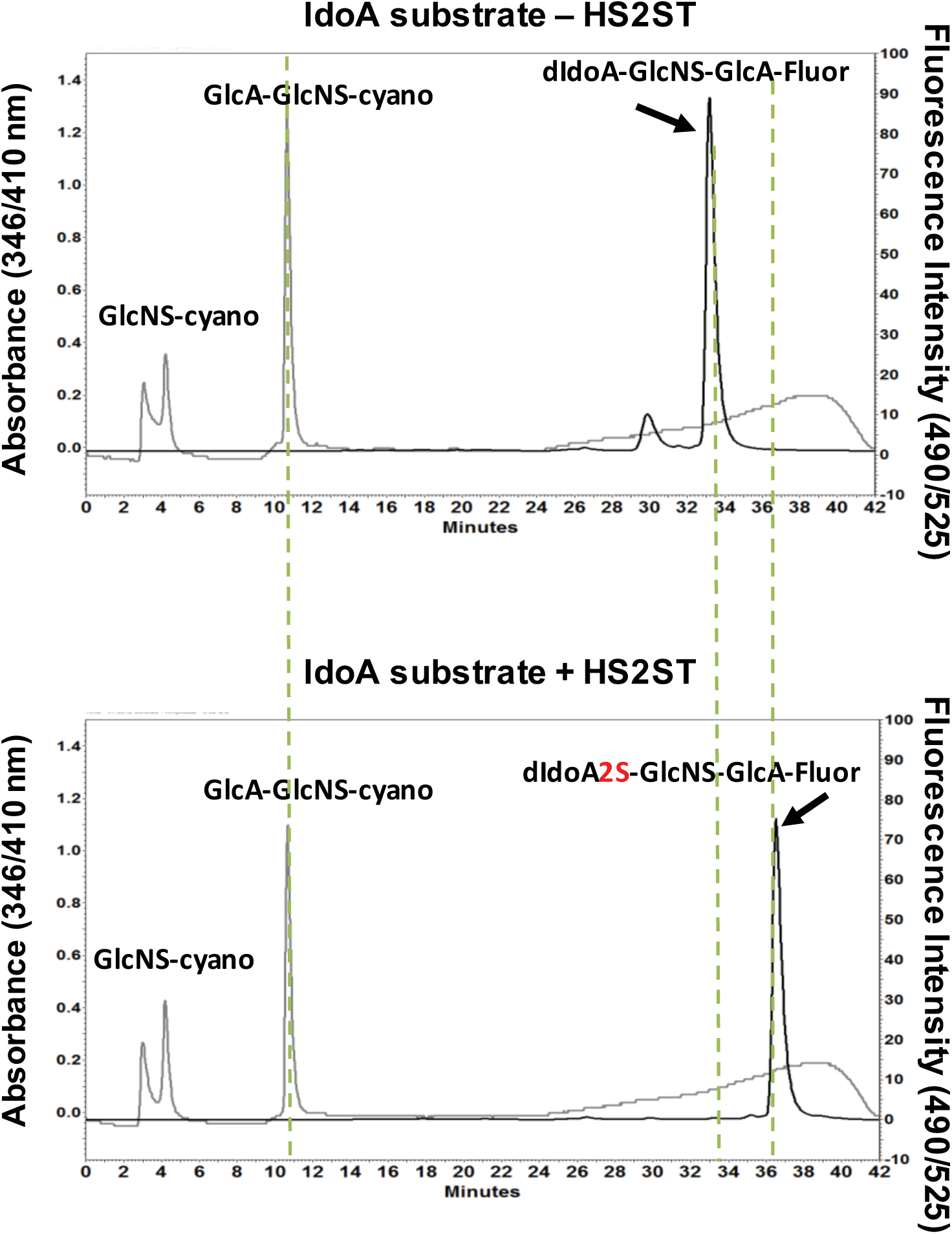
HPLC analysis of sulphated and non-sulphated fluorescent polysaccharide substrate. HPLC separation of cyanoacetamide or fluorescein-labelled saccharides obtained from heparitinase digestion of GlcNS-GlcA-GlcNS-IdoA-GlcNS-GlcA-Fluorescein HS2ST substrate. Elution profiles of digested polysaccharide after anion exchange chromatography are shown. The non-sulphated IdoA-containing hexameric substrate (eluting at ~34 min, top) and the 2-*O*-sulphated product (eluting at ~37 min, bottom) were confirmed by comparison of the different peaks in the fluorescence spectra (dashed lines), with the later eluting sulphated product highlighted in red. dIdoA refers to the double bond formed by β-elimination between C4 and C5 in the IdoA and 2-*O*-IdoA oligosaccharides.

**Supplementary Figure 5.**
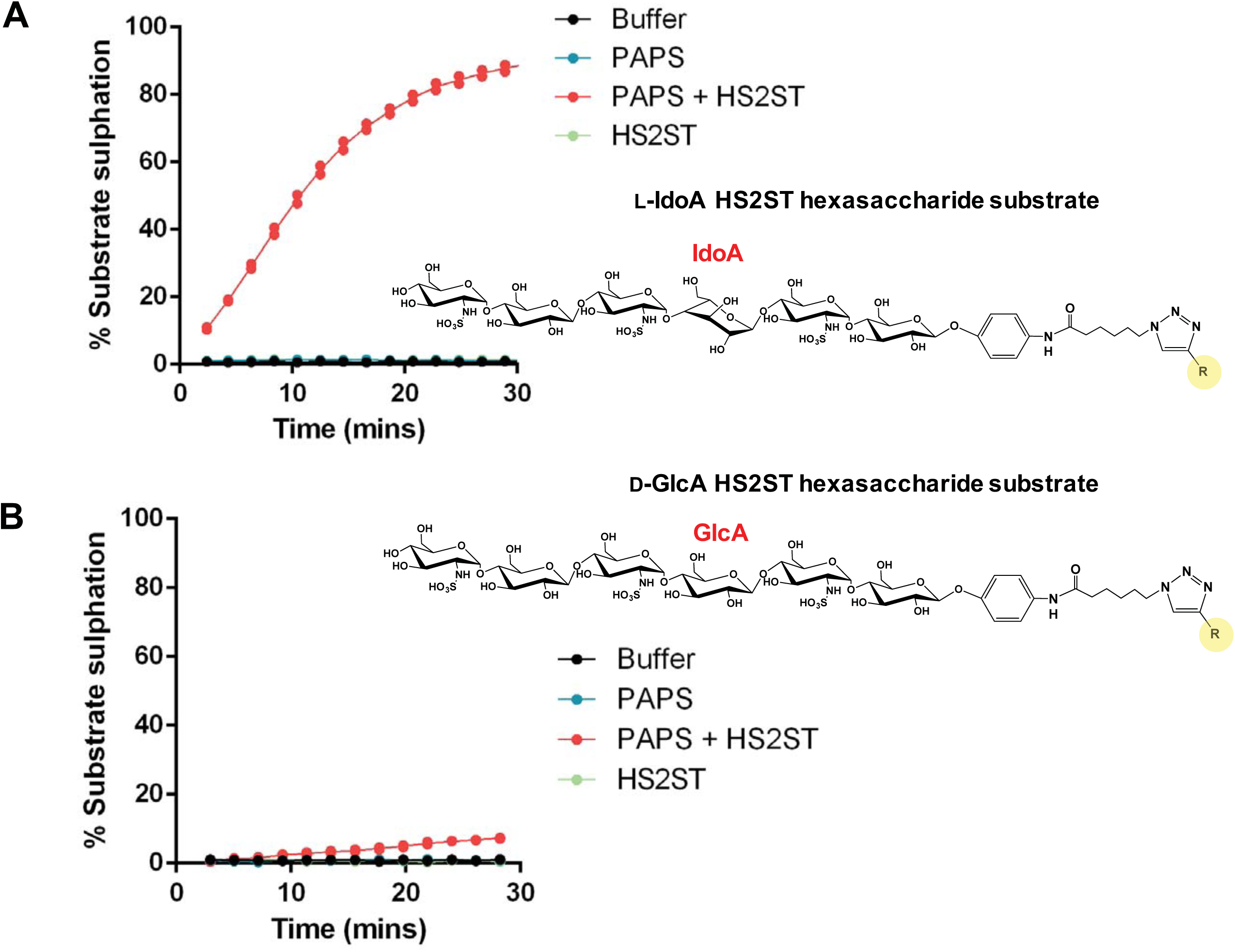
HS2ST glycan residue substrate-specificity analysis. Efficient sulphation of a hexasaccharide substrate by HS2ST requires an L-IdoA residue at the appropriate position in the oligosaccharide. Direct microfluidic sulphotransferase assays demonstrating time-dependent sulphation of the fluorescein-tagged hexasaccharide substrate containing either **(A)** L-IdoA or **(B)** D-GlcA residue at the third residue from the fluorescein-conjugated (reducing) end. R=fluorescein. The IdoA or GlcA residues are indicated in red.

**Supplementary Figure 6.**
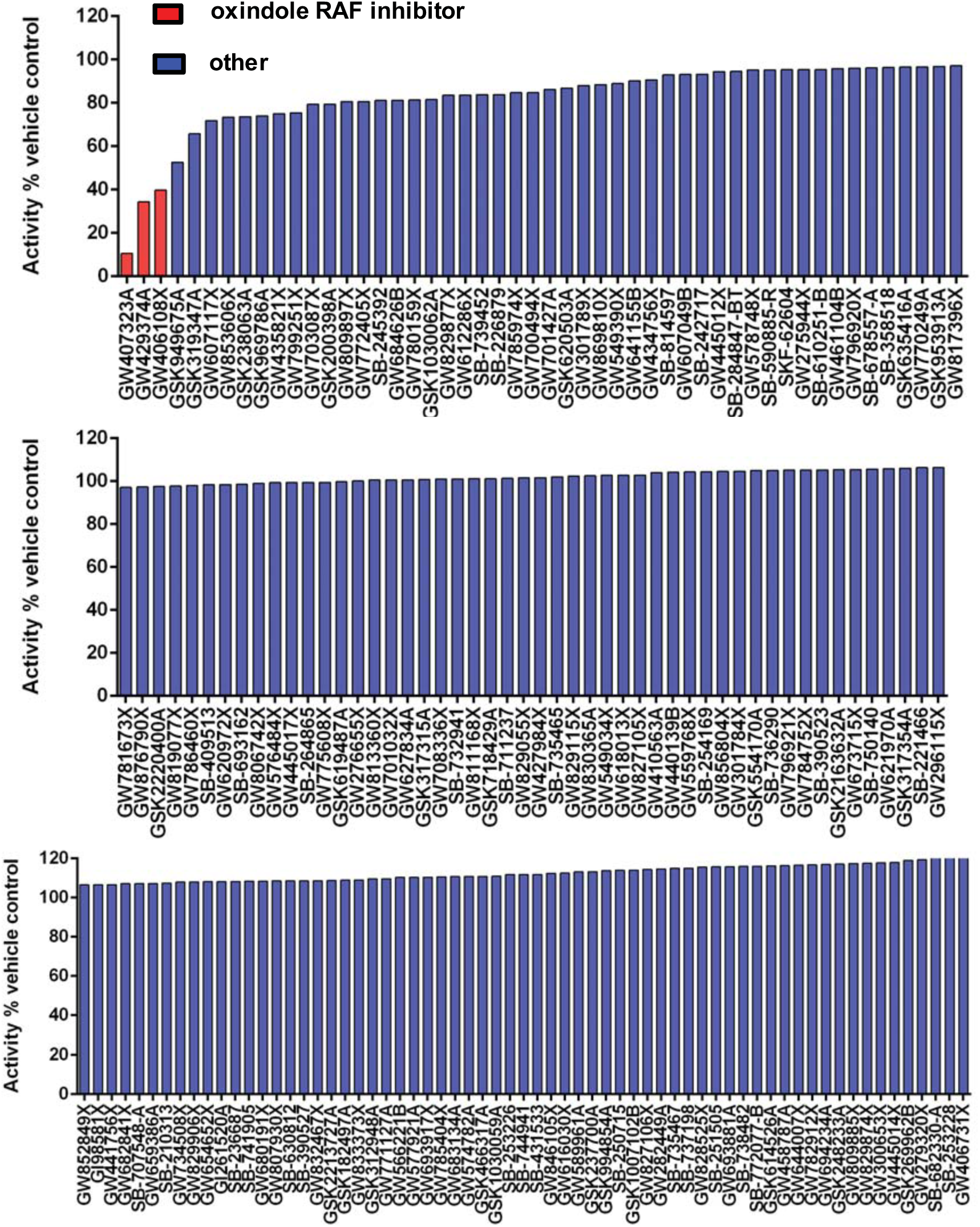
HS2ST enzymatic PKIS compound screen. Inhibition of HS2ST catalytic activity by selected PKIS members. Data are presented as HS2ST activity relative to DMSO control, assayed in duplicate. The most notable ‘hit’ inhibitors from the oxindole chemical class are shaded in red.

**Supplementary Figure 7.**
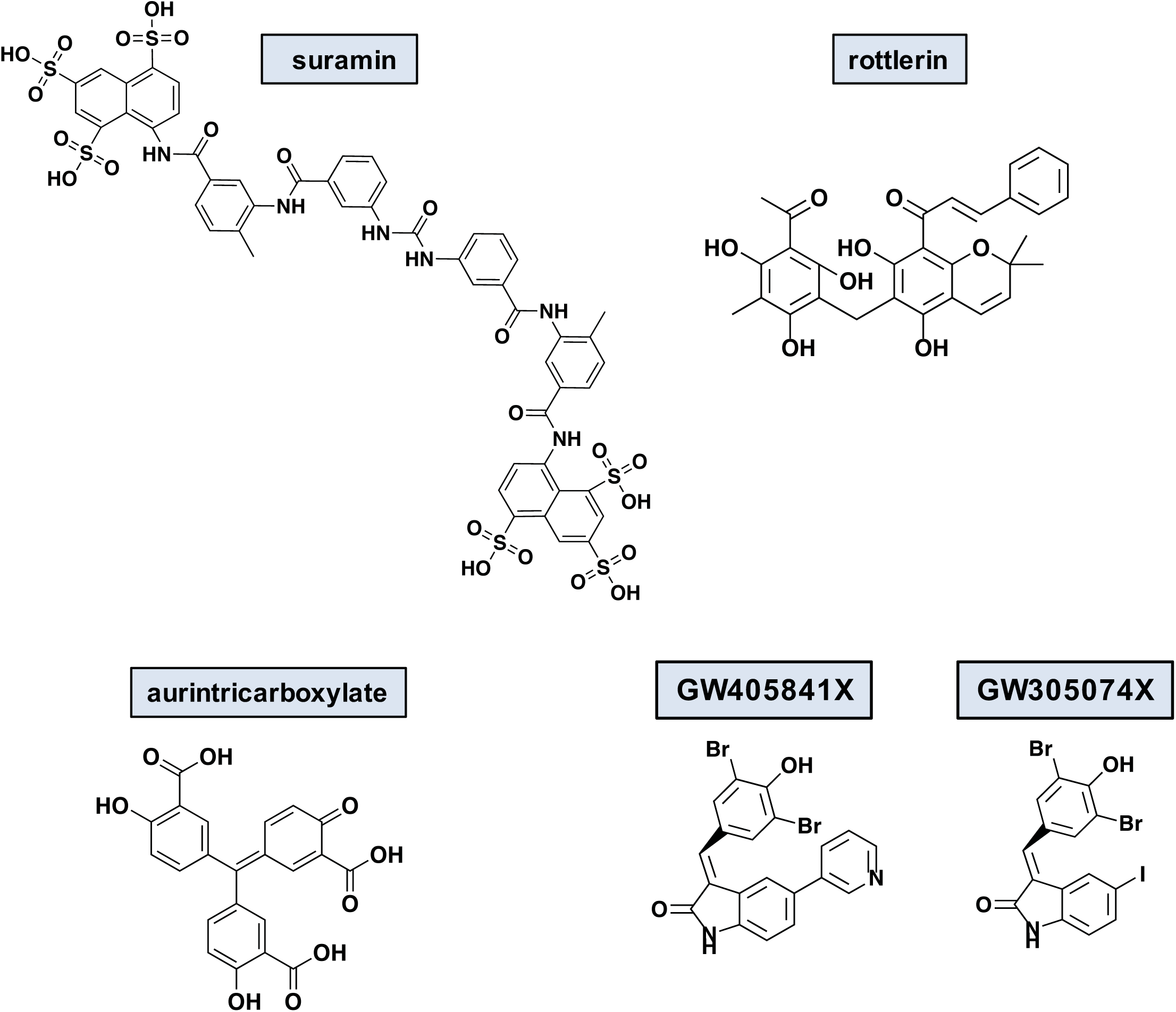
Chemical structures of HS2ST inhibitory ligands. Chemical structures of surmain, rottlerin, aurintricarboxylic acid and selected PKIS compounds.

**Supplementary Figure 8.**
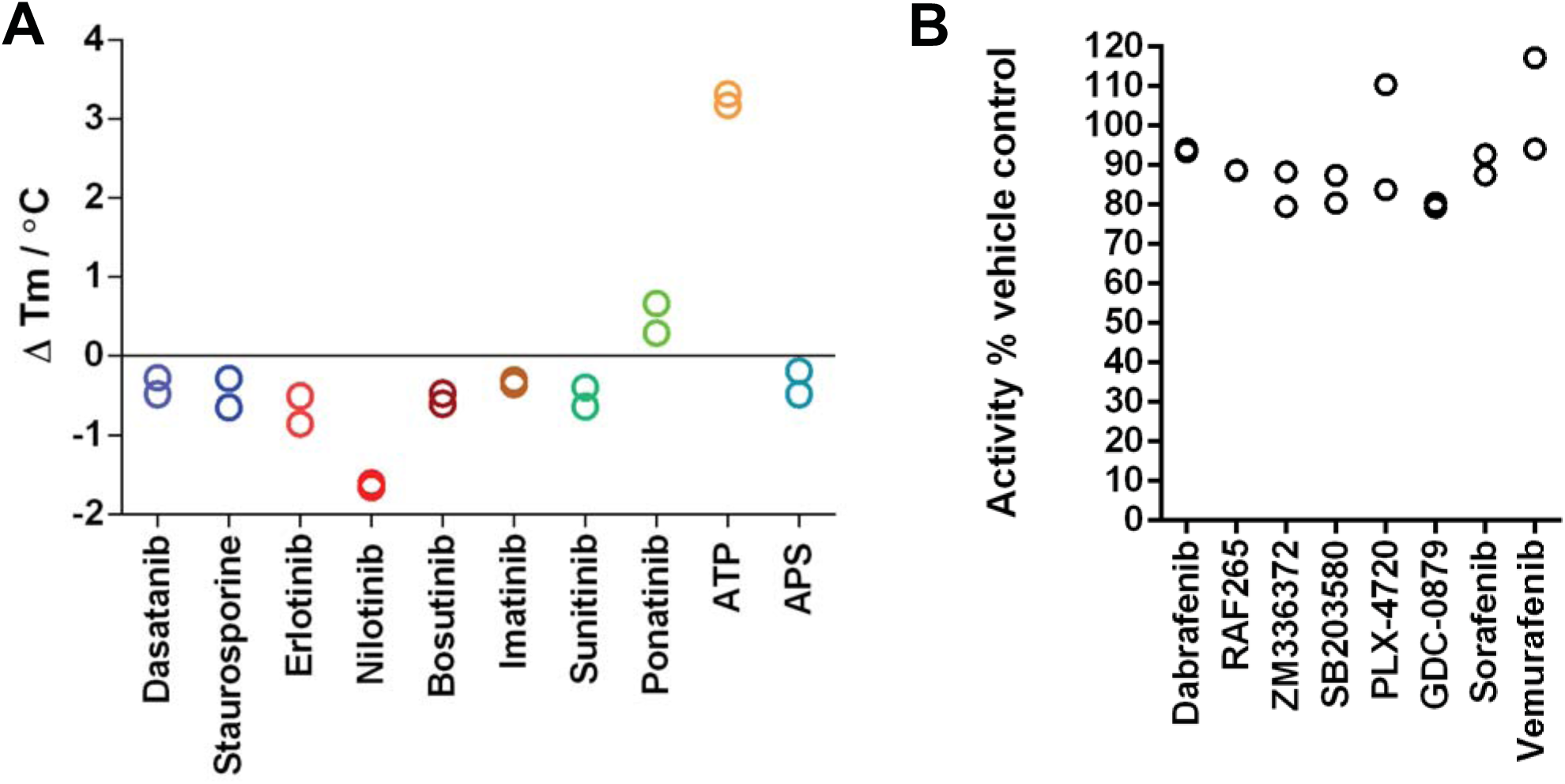
Lack of HS2ST inhibition by various kinase inhibitors. DSF screening (left panel) or enzyme-based inhibitor assay (right panel) evaluating staurosporine, FDA-approved kinase inhibitors and several chemically-distinct RAF kinase inhibitors.

## ABBREVIATIONS

DSF: Differential Scanning Fluorimetry
GlcA: β-D-glucouronate
HS2ST: heparan sulphate 2-*O*-sulphotransferase
IdoA: α-L-iduronate
PAPS: (Adenosine 3′-phosphate 5′-phosphosulphate
PKIS: Published Kinase Inhibitor Set
RAF: Rapidly Accelerated Fibrosarcoma
TSA: Thermostability Assay

